# Immuno-*moodulin*: a new anxiogenic factor produced by autoimmune-prone T cells

**DOI:** 10.1101/796664

**Authors:** Giuseppa Piras, Lorenza Rattazzi, Nikolaos Paschalidis, Silvia Oggero, Giulio Berti, Masahiro Ono, Fabio Bellia, Claudio D’Addario, Bernardo Dell’Osso, Carmine Maria Pariante, Mauro Perretti, Fulvio D’Acquisto

**Author notes:** Correspondence to: Fulvio D’Acquisto, Centre for Biochemical Pharmacology, The William Harvey Research Institute, Barts and the London School of Medicine, Charterhouse Square, London, EC1M 6BQ, United Kingdom. Health Science Research Centre, Department of Life Science, University of Roehampton, London, SW15 4JD, UK. Molecular and Cellular Immunology department, Great Ormond Street Institute of Child Health, UCL, 30 Guilford Street, London WC1N 1EH. Cellular Immunology Laboratory, Center for Basic Research, Biomedical Research Foundation of the Academy of Athens, 11527 Athens, Greece (Nikolaos Paschalidis).

## Abstract

Patients suffering from autoimmune diseases are more susceptible to mental disorders yet, the existence of specific cellular and molecular mechanisms behind the co-morbidity of these pathologies is far from being fully elucidated. By generating transgenic mice overexpressing Annexin-A1 exclusively in T cells to study its impact in models of autoimmune diseases, we made the unpredicted observation of an increased level of anxiety. Gene microarray of Annexin-A1 CD4^+^ T cells identified a novel anxiogenic factor, a small protein of approximately 21kDa encoded by the gene 2610019F03Rik which we named Immuno-*mood*ulin. Neutralizing antibodies against Immuno-*mood*ulin reverted the behavioral phenotype of Annexin-A1 transgenic mice and lowered the basal levels of anxiety in wild type mice; moreover, we also found that patients suffering from obsessive compulsive disorders show high levels of I*mood* in their peripheral mononuclear cells. We thus identify this protein as a novel peripheral determinant that modulates anxiety behavior. Therapies targeting Immuno-*mood*ulin may lead to a new type of treatment for mental disorders through regulation of the functions of the immune system, rather than directly acting on the nervous system.

---

The challenging life of patients diagnosed with autoimmune diseases is often further impoverished by the emergence of mental disorders as a major co-morbidity ^1,2^. For instance, ∼40% of patients suffering from multiple sclerosis have attempted suicide ^3,4^ while more than 30% of those affected by autoimmune hepatitis suffer from schizophrenia ^5,6^. Most strikingly, immunomodulatory therapies for the treatment of autoimmune conditions might aggravate the emergence of these problems ^7^ thus presenting both clinicians and patients with a paradoxical dilemma: the physical symptoms of autoimmunity might be effectively improved at the expenses of a worsening of the emotional state and wellbeing. This is for instance the case of interferon beta (IFN-β) that is currently used as an effective treatment for multiple sclerosis but its use is limited by the increased incidence of suicidal thoughts in a significant proportion of patients ^8^. Although some studies have investigated the functional cross-talk between the brain and the immune system^9,10^, it is still not clear how one system influences the other and if there is a common root or determinant for the emergence of mental disorders in autoimmune conditions.

Annexin-A1 (AnxA1) is an endogenous modulator of a variety of physiological and pathological processes ranging from inflammation ^11,12^ to autoimmunity ^13,14^ and cancer ^15-17^. As with many other multifunctional mediators, AnxA1 plays a homeostatic role in the immune system as it can exert both positive and negative functions depending on the contexts. In the context of T cells, studies have indeed provided contrasting and opposite results showing that it can act as both a positive ^18-27^ and a negative modulator of T cell activation ^27-30^. All these studies have been done using either exogenously administered recombinant AnxA1 (and its mimetic) or AnxA1-deficient mice where the protein is absent in every immune cell. Aiming to address these controversies, we have generated a T cell specific AnxA1 transgenic mouse colony (AnxA1^tg^) to address the biology of this mediator.

In this paper, analysis of AnxA1^tg^ response in experimental model of autoimmunity showed exacerbated signs of inflammation. However, besides their autoimmune-prone phenotype, AnxA1^tg^ mice displayed an unexpected high level of compulsive digging that was noticeable even in their home cage and at basal physiological settings e.g. in mice that were not subjected to any inflammatory condition. Further exploration on the molecular mechanisms behind this phenotype of anxiety led us to the discovery of a new homeostatic modulator that we named Immuno-*mood*ulin (I*mood*) - because of its discovery in T cells. Immuno-*mood*ulin levels in mice affect their intrinsic basal level of anxiety. In addition, we found that patients suffering from OCD present high levels of I*mood* in their peripheral blood mononuclear cells. The results of this study expand our knowledge of the complex interactions and intersection between the immune and the central nervous system.

## Materials and Methods

### Generation of T cell-specific AnxA1^tg^ mice

To generate the VACD2 AnxA1 FLAG transgenic mice, murine AnxA1 gene was extracted, amplified and tagged with the FLAG epitope. The gene was cloned in TOPO pcDNA3.1 vector for verification of its expression *in vitro* and finally subcloned in the VACD2 vector ^31^ for T cell specific expression in the mouse. Finally, the VACD2 AnxA1 FLAG construct was modified and purified for the pronuclear microinjection into the mouse genome.

### Subjects

20 OCD outpatients of either gender and any age, treated and followed up at the OCD tertiary outpatient Clinic of the University Department of Psychiatry of Milan, Policlinico Hospital, were included in the study. Diagnoses were assessed by the administration of a semi-structured interview based on DSM-5 criteria (SCID 5 research version, RV). In case of psychiatric comorbidity, OCD had to be the primary disorder, causing the most significant distress and dysfunction and providing the primary motivation to seek treatment. Patients were excluded from the study if they had recent or current alcohol or substance abuse (last 3 months), as well as medical conditions including autoimmune diseases, due to their potential influence over gene expression. For the same reason, lifetime history of trauma (according to DSM-5), as well as the current presence of relevant psychological stress, were considered exclusion criteria. Clinical assessment included the collection of the following demographic and clinical variables: gender, age, age at onset, and current pharmacological treatment. In addition, illness severity was measured through the Yale-Brown Obsessive-Compulsive Scale. Patients had maintained their pharmacological treatment stable for at least one month in order to be enrolled in the study and on the day of collection of blood. Control subjects (n = 20) were volunteers matched for gender, age and ethnicity, with no psychiatric diagnosis as determined by the SCID 5 and no positive family history for major psychiatric disorders in the first-degree relatives (as assessed by the Family Interview for Genetic Studies). Blood collection was performed between 2 and 4 pm from not fasting donors. PBMCs were separated by density gradient using the Lympholyte-H kit as recommended by the manufacturer (Cedarlane Laboratories, Canada). All subjects had given their written informed consent to participate to the study, which included the use of personal and clinical data as well as blood drawing for genotyping and methylation analysis. The study protocol had been previously approved by the local Ethics Committee.

## Results

### Generation and phenotypic characterization of T cell specific AnxA1^tg^ mice

We generated T cell specific transgenic mice through pronuclear injection of VACD2 AnxA1 FLAG construct in 129 FVB mice (**Supplementary Figure 1**). Both the two founders and their litters showed no gross sign of disease. Following backcrossing onto C57BL/6 background and intercross to generate mice with the transgene on both alleles, we noticed that the female litters from one of the two transgenic founders presented an unusual high incidence (almost 80%) of maternal cannibalism. This was successfully controlled by administering perphenazine (0.025 mg/ml in drinking water) during pregnancy as it has been previously reported for other autoimmune prone and highly anxious mouse strains such as the DBA/2J ^32^. Rescued newborn pups from this line (called ‘the red line’) had often to be raised by C57BL/6 foster mothers to avoid losing the colony. The combined administration of perphenazine and fostering care, made this line challenging to use. Therefore, unless otherwise stated, all the experiments were carried out using the litters of the other founder. Analysis of the immune repertoire of AnxA1^tg^ mice from both founders showed no significant differences in lineage commitment towards CD4^+^ or CD8^+^ cells in all lymphoid organs (**Supplementary Figure 2A**). To assess if AnxA1 overexpression in T cells would have any effects on the cellularity of the lymphoid organs we measured the total cell number of cells present in lymph nodes, spleen and thymus of AnxA1^tg^ mice compared to control. As shown in **Supplementary Figure 2B**, no significant changes were found except in in lymph nodes where the cellularity was significantly higher (by about 60%) in AnxA1^tg^ mice compared to control.

### Increased T cell activation and autoimmunity in AnxA1^tg^ mice

Consistent with our previous reports ^19,20,22,23^, AnxA1^tg^ T cells showed a clear pro-inflammatory phenotype as evidenced by lower threshold of CD25 and CD69 upregulation (**Figure 1A**) and increased production of IL-2 following anti-CD3 plus anti-CD28 (anti-CD3/CD28) stimulation (**Figure 1B**). *In vivo*, AnxA1^tg^ mice showed an exacerbated inflammatory response in the MOG_35-55_-induced experimental autoimmune encephalomyelitis (EAE) (**Figure 1C**) as evidenced by the exacerbated severity of the clinical score and increased weight loss after the onset of the disease (day 12) and larger inflammatory infiltrate of the spinal cord in AnxA1^tg^ mice compared to wild type. No differences were observed in terms of disease incidence between control and AnxA1^tg^ mice (**Table in Figure 1C**). The same exacerbated inflammatory response was observed and confirmed in the red line mice (**Supplementary Figure 3A**). To expand and confirm these findings in another model of autoimmune inflammation, we subjected AnxA1^tg^ mice to an experimental model of systemic lupus erythematosus ^33^. As shown in **Figure 1D** (right panel), injection of pristane to AnxA1^tg^ mice provoked a significant weight loss over a period of 35 days while control mice gained about 30% of their initial weight over the same period. As the disease progresses, a number of organs, including the spleen and the lung, becomes enlarged or accumulate inflammatory infiltrate ^34,35^. As shown in **Supplementary Figure 4A**, AnxA1^tg^ mice showed increased splenomegaly (spleen weight 0.130± 0.010 gr in wild type mice vs 0.281± 0.011 gr in AnxA1^tg^; n=5 mice, p<0.0001) with typical pathological features such as enlarged B-cell follicles and infiltration of oil droplets compared to control wild type mice. The lungs of AnxA1^tg^ mice also showed increased areas of focal hemorrhages together with severe mixed inflammatory infiltrate compared to control (**Supplementary Figure 4B)**. Consistent with these results, ∼ 30% of AnxA1^tg^ mice died during the 35-day treatment while control animals showed a 100% survival (**Figure 1D**; left panel).

**Figure 1.**
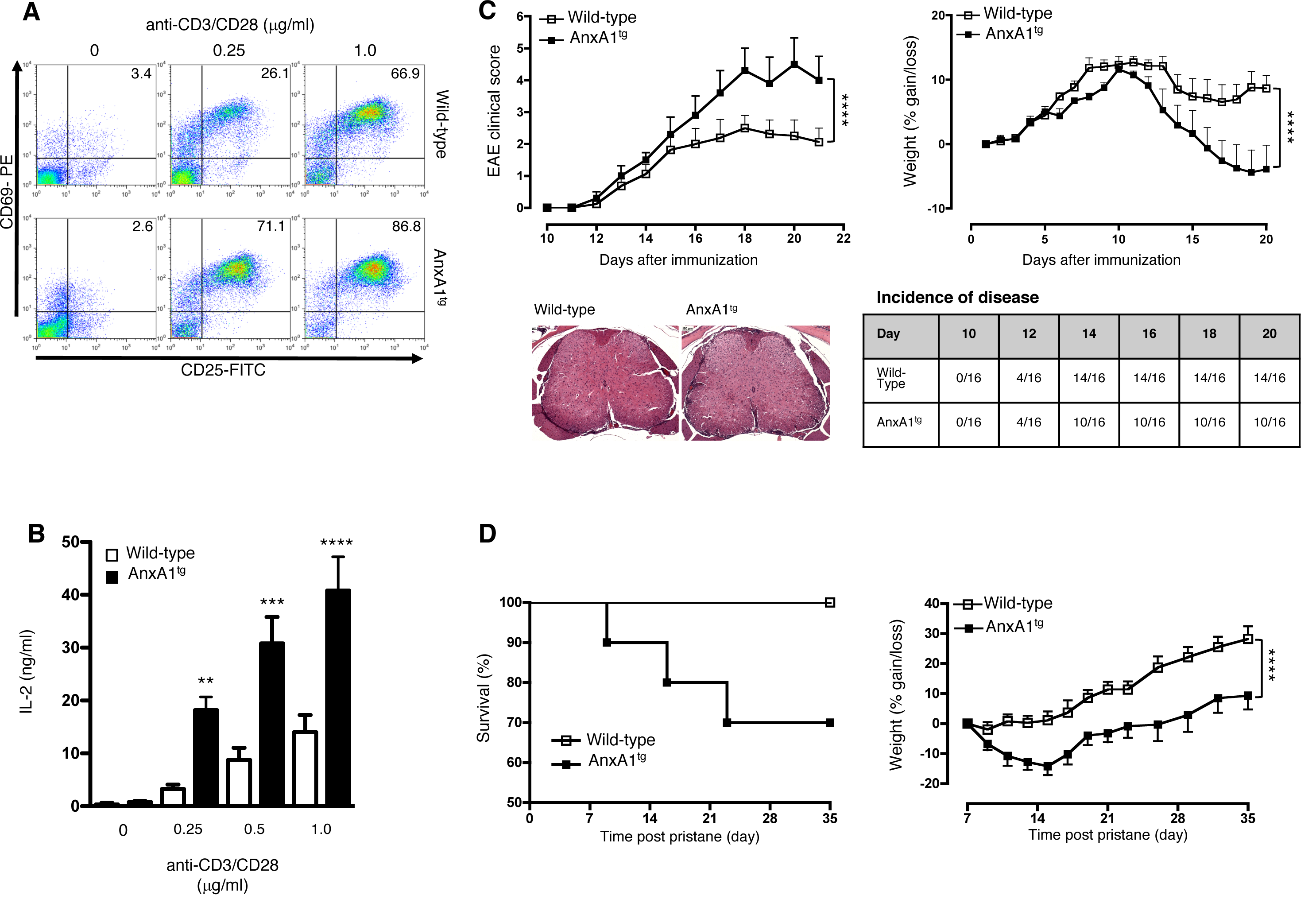
Autoimmune-prone phenotype of T-cell specific AnxA1^tg^ mice. (**A**) T cells from control and AnxA1^tg^ mice were stimulated with the indicated concentration of plate-bound anti-CD3/CD28 for 16-18 hrs and then stained with anti-CD25 and anti-CD69 and analyzed by FACS. The numbers in the plot show the percentages of CD69 and CD25 double positive cells. Results are representative of n=12-18 experiments with similar results. (**B**) T cells from control and AnxA1^tg^ mice were stimulated with the indicated concentration of plate-bound anti-CD3/CD28 for 24-30 hr and the supernatants used to measure the levels of IL-2. The bars show means ± SEM n=5 separate mice and are representative of five experiments with similar results. Statistical significance between control and AnxA1^tg^ cells was determined by two-way ANOVA followed by Bonferroni multiple-comparison test ** p<0.01; *** p<0.001; ****p<0.0001. (**C**) Control and AnxA1^tg^ mice were immunized with MOG_35-55_ and CFA and monitored daily for clinical signs of EAE (top left panel) or weight gain/loss (top right panel) for 23 days. Results show means ± SEM of n=8 mice per group and are representative of seven experiments with similar results. **** p<0.0001 (two-way ANOVA followed by Bonferroni multiple-comparison test) indicates significant values compared to wild type control mice. The spinal cord sections showed in the bottom left panels were obtained at day 18 and stained with hematoxylin and eosin as described in Materials and Methods. The table in the bottom right corner shows the number of mice showing a score of 2 at different times during the development of the EAE. (**D**) Control and AnxA1^tg^ mice received an intraperitoneal injection of pristane to induce a lupus-like disease and were monitored daily for survival (left panel) or weight gain/loss (right panel) for 35 days. Results show means ± SEM of n=10 mice per group and are representative of three experiments with similar results. *** p<0.001 (two-way ANOVA followed by Bonferroni multiple-comparison test) indicates significant values compared to wild type control mice.

### Selective accumulation of Th1/Th17 double positive cells in the inflamed tissue of AnxA1^tg^ mice

To further examine the activation state of AnxA1^tg^ T cells in the EAE mice, we investigated the effector phenotype of these cells at day 9 (onset of the disease) and day 16 (peak of the disease). This allowed us to distinguish the effects of AnxA1 overexpression during the priming phase occurring in the draining lymph nodes at day 9 or the differentiation phase occurring within the spinal cord at day 16 ^36^. Comparison of the total cell number of CD45^+^ leukocytes in either lymph node or spinal cord of AnxA1^tg^ and wild type mice showed no significant differences (**Figure 2A and B, bottom panels**). Day 9 comparison of the draining lymph nodes of control and AnxA1^tg^ mice showed a significant increase in the number of CD4^+^ T cells (by about 85%) in the latter group, but no difference in the percentage of IL-17^+^/IFN-γ^+^ or IL-17^+^/GM-CSF^+^ double-positive or single-positive T cells (**Figure 2A**). At day 16, the number of CD4^+^ T cells recovered from the spinal cord of was ∼3-fold higher in AnxA1^tg^ compared to control but in this case we observed an increase in the percentages of IFN-γ^+^/IL-17^+^ (1.9 ± 0.12% in wild type vs 4.5 ± 0.21% in AnxA1^tg^; p<0.0001) or GM-CSF^+^/IL-17^+^ (4.8 ± 0.19% in wild type vs 10.4 ± 0.45% in AnxA1^tg^; p<0.0001) pathogenic ^37^ double-positive T cells in the former compared to the latter (**Figure 2B**). Fate mapping reporter studies for Th17 cells in this model of autoimmune inflammation have shown that these double-positive cells represent a transition phase during the ‘conversion’ of IL-17 single-positive into IFN-γ or GM-CSF double-positive T cells at the site of inflammation ^38^. Consistent with these findings, AnxA1^tg^ mice show a higher percentage of IL-17^+^ cells and a corresponded reduced percentage of IFN-γ (about 10% in AnxA1^tg^ *vs* 18 % in control) and GM-CSF (about 16 % in AnxA1^tg^ *vs* 20% in control) single-positive cells (**Figure 2B**).

**Figure 2.**
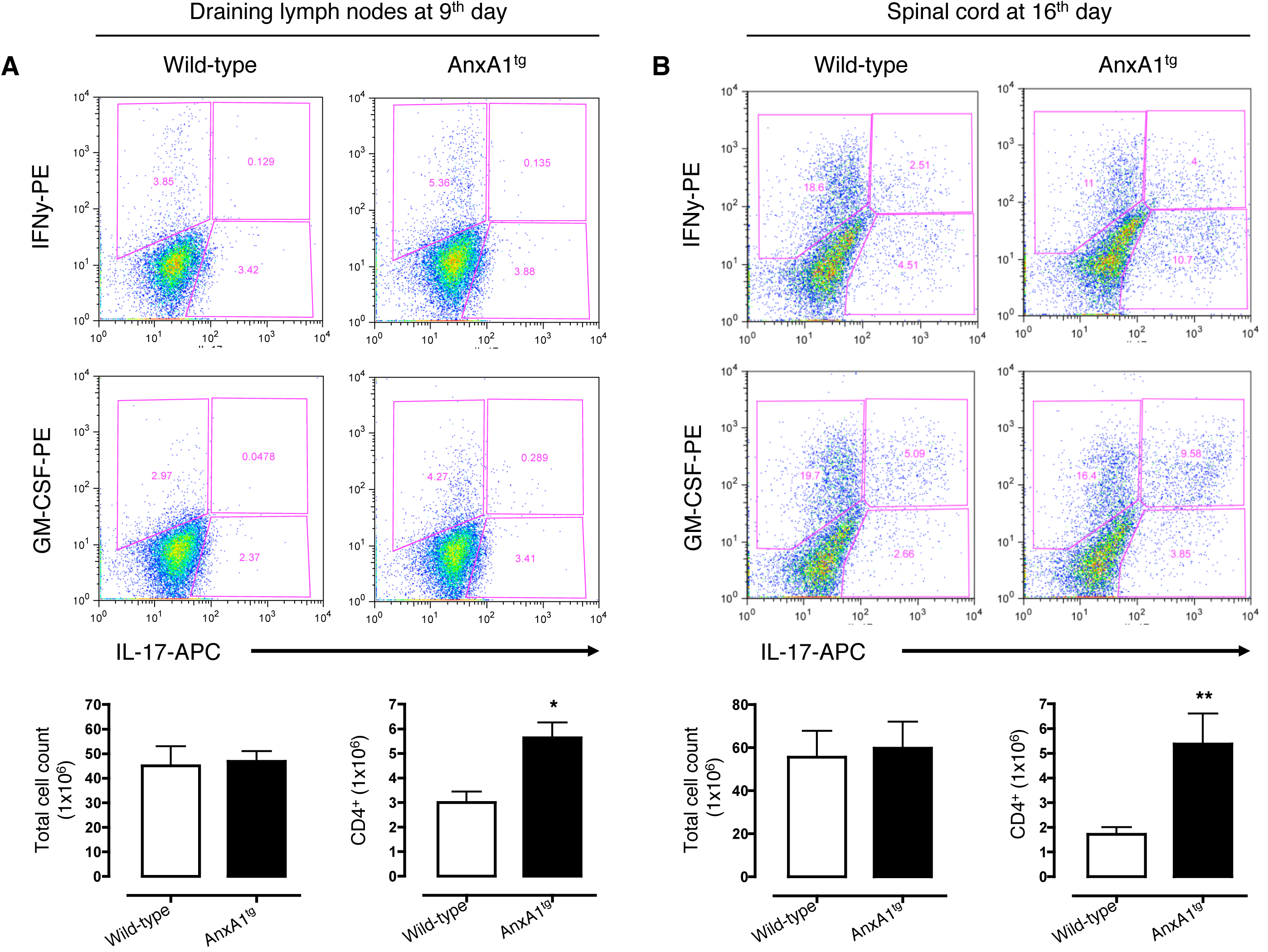
Phenotype of effector T cells recovered from AnxA1^tg^ mice subjected to MOG_35-55_-induced EAE. Representative dot plots showing intracellular staining for IL-17 and IFN-γ (top panels) or IL-17 and GM-CSF (bottom panels) of CD4^+^ T cells present in the draining lymph nodes (**A**) or spinal cords (**B**) of wild type and AnxA1^tg^ T cells sacrificed at the indicated time after MOG_35-55_ immunization. The bar graphs at the bottom show the total cell number of CD45^+^ mononuclear cells recovered after Ficoll (draining lymph nodes) or Percoll (spinal cord) purification as described in the Supplementary Materials and Methods. The bar graphs referring to CD4^+^ T cells show the number of these cells present in the same tissues. Values are expressed as means ± SEM of n=6 mice for each group and are representative of three experiments with similar results. *p<0.05; **p<0.01 (Student’s *t*-test) indicate significant values compared to wild type control mice.

### Increased basal anxiety-like behavior in AnxA1^tg^ mice

Direct observation of AnxA1^tg^ mice in their cage revealed an altered behavior typified by an increased tendency to compulsive digging (compare **Movie 1**, wild type mice and **Movie 2**, AnxA1^tg^ mice). Similar behavior was observed in the litters from the other founder (red line mice; compare **Movie 3** of wild type mice with **Movie 4** of AnxA1^tg^ mice). Interestingly, the litter of red line mice also showed an increased level of hetero-barbering (plucking of fur from cage-mates; **Movie 5**) - a behavior that has been proposed to represent a model of human trichotillomania and obsessive-compulsive spectrum disorders^39^. To thoroughly measure this heightened anxious behavior, we used a battery of classical tests for anxiety behavior. The marble-burying test is used to measure digging ^40^ and when applied to AnxA1^tg^ we quantified a significant increase in the number of buried marbles by the mice, which spent approximately double the time on this activity (**Figure 3A**). In the light and dark shuttle box test ^41^ AnxA1^tg^ mice spent in the lit area than the wild type counterpart, with a marked attenuation of numbers of crossings between the two compartments (**Figure 3B**). Significant behavioral differences were observed also in the climbing test-an experimental paradigm to measure vertical activity ^42^: AnxA1^tg^ mice showed an increased latency to the first climb and a significant reduction in the time spent on this activity (**Figure 3C**). All these changes were not secondary to a general impairment of locomotor activity as AnxA1^tg^ mice showed no difference in the number of i) square crossed or ii) rearing as quantified in the open field test ^43^ when compared to wild type animals (**Figure 3D**). This heightened anxiety-like behavior was confirmed in the litters of the red line mice (**Supplementary Figure 3B**).

**Figure 3.**
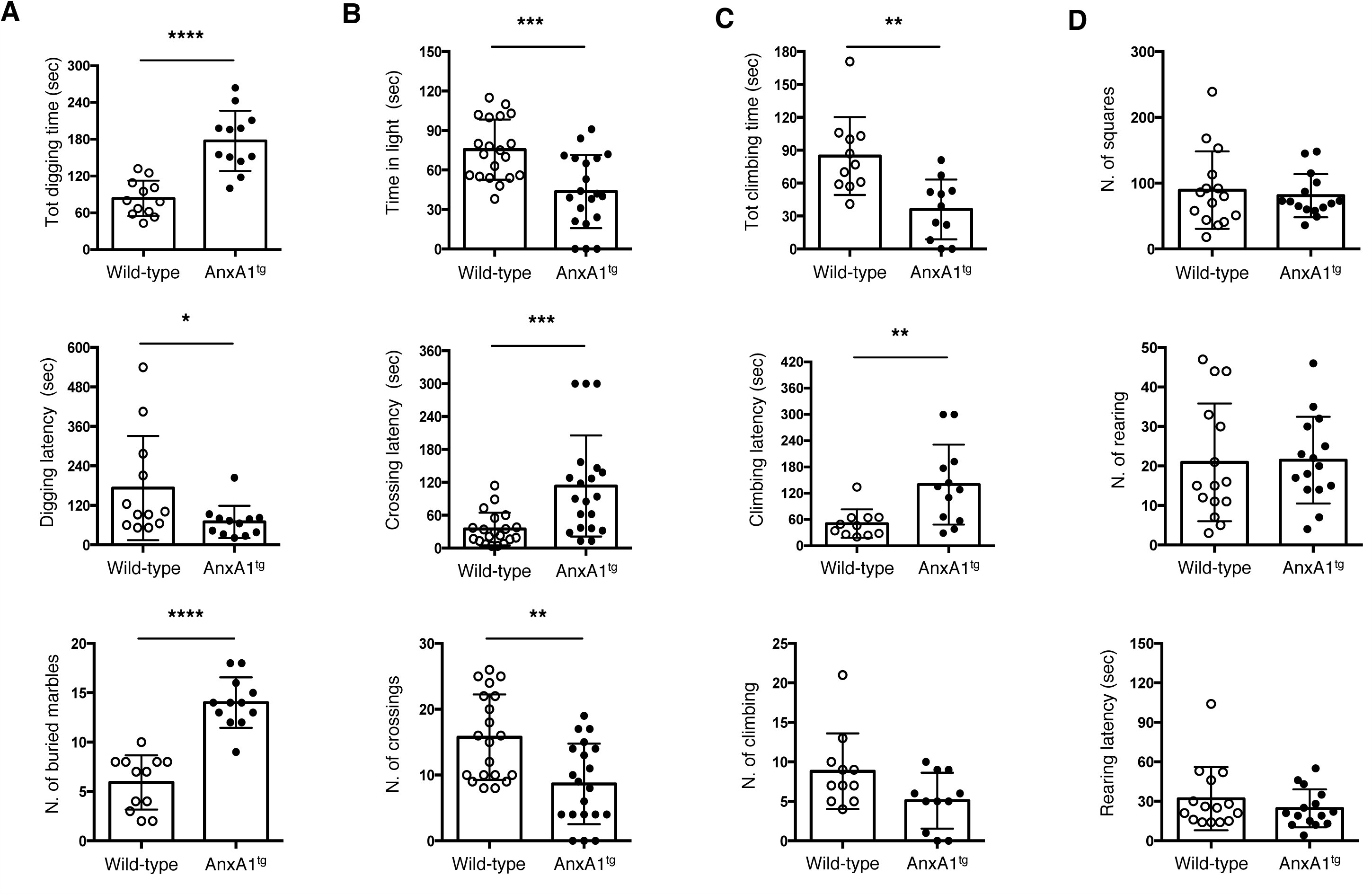
Increased signs of anxiety-like behavior in T cell-specific AnxA1^g^ mice. (**A**) The bar graphs show the total duration (seconds) of digging, the latency (seconds) to the first digging bout and the total number of buried marbles during a 10-minute trial. (**B**) The bar graphs show the total time (seconds) spent in the lit area, latency (seconds) to first cross to the dark chamber and total number of crossings during a 5-minute trial. (**C**) The bar graphs show, the total time (seconds) spent on the climbing mesh, the latency (seconds) to the first climb and the number of climbing events during a 5-minute trial. (**D**) The bar graphs show total number of squares crossed, the number of rears and the latency (seconds) to the first rear during a 5-minute session. Values are expressed as means ± SEM of n=12 - 20 mice and are representative of four different experiments with similar results. * p<0.05, **p<0.01, ***p<0.001, ****p<0.0001 indicate significant values compared to wild type control mice (Mann–Whitney U-test).

### Gene fingerprint of AnxA1^tg^ whole brain reveals an increased expression of anxiety-related genes

CD4^+^ T cells exert a homeostatic control over a number of functions of the brain including learning, memory and anxiety-like behavior ^44^. In a previous study we have shown that CD4^+^ rather than CD8^+^ T cells revert the increased anxiety-like behavior observed in Rag-1^-/-^ immunodeficient mice. Most interestingly, we have shown that the modulatory effects exerted by CD4^+^ T cells on behavior were ‘mirrored’ by specific changes in the gene fingerprint of the whole brain ^45^. Thus, for instance, S100a10, a known biomarker for anxiety-like behavior, was up-regulated in RAG-1^-/-^ and down-regulated to wild type level in RAG-1^-/-^ mice where CD4^+^ T cells artificially re-introduced (RAG-1^-/-^/OT-II TCR transgenic mice). In light of these premises, we queried if AnxA1^tg^ mice would also show changes in their brain gene fingerprint and performed a comparative analysis of gene expression of the whole brain using microarray analysis. The results showed significant differences in the level of expression of 15 genes of which 8 were unregulated and 7 downregulated in the whole brains of AnxA1^tg^ mice in comparison to those of wild type mice (**Figure 4A** and **Supplementary Table 1**). Among these, several were associated with emotional disorders including alcoholism and anxiety such as erythroid differentiation regulator 1 (*Erdr1)* ^46^ and gamma-aminobutyric acid receptor subunit alpha-2 (*Gabra2)* ^*47*^ (**Figure 4B**) RT-PCR analysis of these genes on a larger number of samples confirmed these results with a down regulation of *Erdr1* by about 78% and an ∼3-fold upregulation of *Gabra 2* (**Figure 4C**). Thus, the transgenic T cells can exert a tonic regulation on a discrete set of genes in the brain, even in the absence of any experimental manipulation.

**Figure 4.**
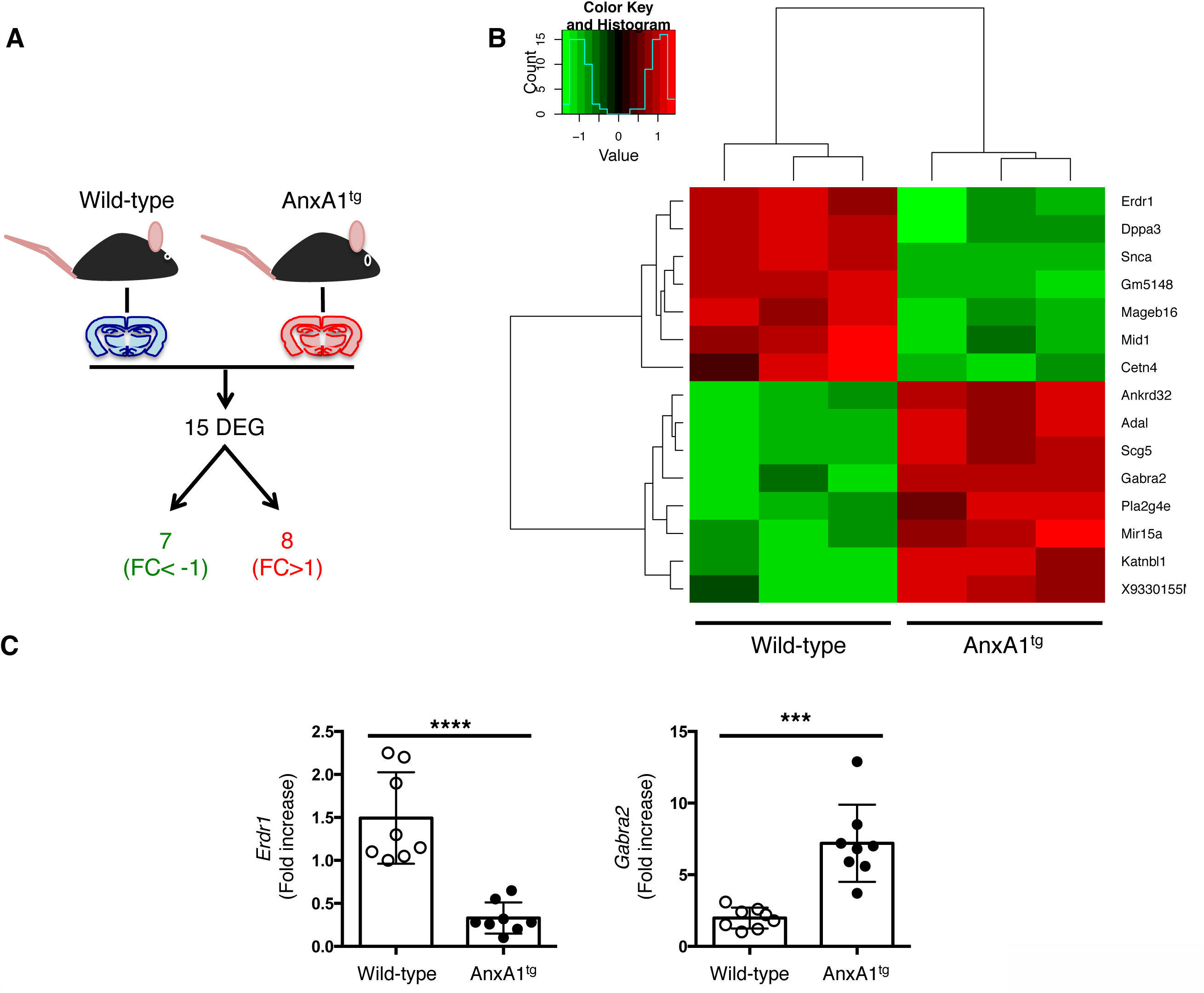
Gene fingerprint of the whole brain of T cell-specific AnxA1^tg^ mice reveals several genes linked to anxiety-like behavior. (**A**) Summary of the gene expression profile of the whole brain of wild type and AnxA1^tg^ mice showing the number of upregulated (FC>1) and downregulated (FC<1) differentially regulated genes (DEG). (**B**) Heatmap analysis on Microarray data of the whole brain of n=3 wild type and AnxA1^tg^ mice. The heatmap analysis used annotated genes only (genes with EntrezID). (**C**) Real time PCR analysis of two genes of interest selected from the microarray. Values are presented as individual data points ± SEM of n=8 mice. ***p<0.001; ****p<0.0001 indicate significant values compared to wild type control mice (Student’s *t*-test).

### AnxA1^tg^ CD4^+^ T cells express high level of a new anxiogenic factor named Immuno-moodulin

To identify the anxiogenic factor produced by AnxA1tg CD4^+^ T cells, we compared gene expression profiles of purified CD4^+^ T cells by microarray. In resting conditions, no statistical differences were observed between AnxA1^tg^ and control CD4^+^ T cells (data not shown). However, in anti-CD3/CD28 stimulated cells, 8 genes were identified to be significantly modulated (**Supplementary Table 2)**. Among those upregulated, AnxA1, interferon-inducible 203 and 2610019F03Rik (also called testis development related protein, *Tdrp* or C8orf42). We were intrigued by the last one as it encoded for a small protein of about 21kDa (like most of the cytokines) that was stored in vesicles (https://www.proteinatlas.org/ENSG00000180190-TDRP/cell) and present in circulation (https://genecards.weizmann.ac.il/v3/cgi-bin/carddisp.pl?gene=TDRP) and therefore decided to investigate its function. As we identified this gene in T cells and hypothesized that would be responsible for the anxious behavioral phenotype of AnxA1^tg^ mice, we named it Immuno-*mood*ulin (I*mood*).

Next, we validated microarray results using immunoblotting and FACS intracellular staining of CD4^+^ T cells using a commercially available polyclonal antibody against I*mood*. We first investigated I*mood* expression in resting or activated T cells from non-transgenic C57BL/6 mice. In basal conditions, about 20% of cells express high levels of I*mood*. Activation of CD4^+^ T cells *via* the TCR caused a clear increase in the number of these cells with a doubling of their number (44%) following the triggering of both signal 1 (anti-CD3) and signal 2 (anti-CD28) (**Figure 5 A**). When we compared the expression of I*mood* in resting wild type and AnxA1^tg^ CD4^+^ T cells (**Figure 5 B**), it was possible to see an increase in the percentages of I*mood*-high cells in AnxA1^tg^ mice compared to wild type (about 55% in AnxA1^tg^ *vs* 35% in wild type) and their median fluorescence intensity (about 15 in AnxA1^tg^ *vs* 7 in wild type). These differences tapered off but remained visible following activation with anti-CD3/CD28: about 80% of AnxA1^tg^ cells expressed I*mood* with a median fluorescence intensity of 26 while about 70% wild type cells showed a median of 18. These results were confirmed at mRNA levels where it was possible to observe both the upregulation of I*mood* mRNA following activation of T cells with anti-CD3/CD28 and an increased expression in AnxA1^tg^ compared to wild type in both resting and stimulating conditions (**Figure 5 C**). We confirmed these findings by western blot. Resting AnxA1^tg^ CD4^+^ T cells showed increased levels of intracellular I*mood* compared to control (Figure 6D, Top panel). In addition, immunoprecipitation of I*mood* from the cell supernatant of wild type and AnxA1^tg^ T cells in both resting and activated conditions showed an increased secretion of this protein in AnxA1^tg^ T cells (**Figure 5 D**, bottom panel).

**Figure 5.**
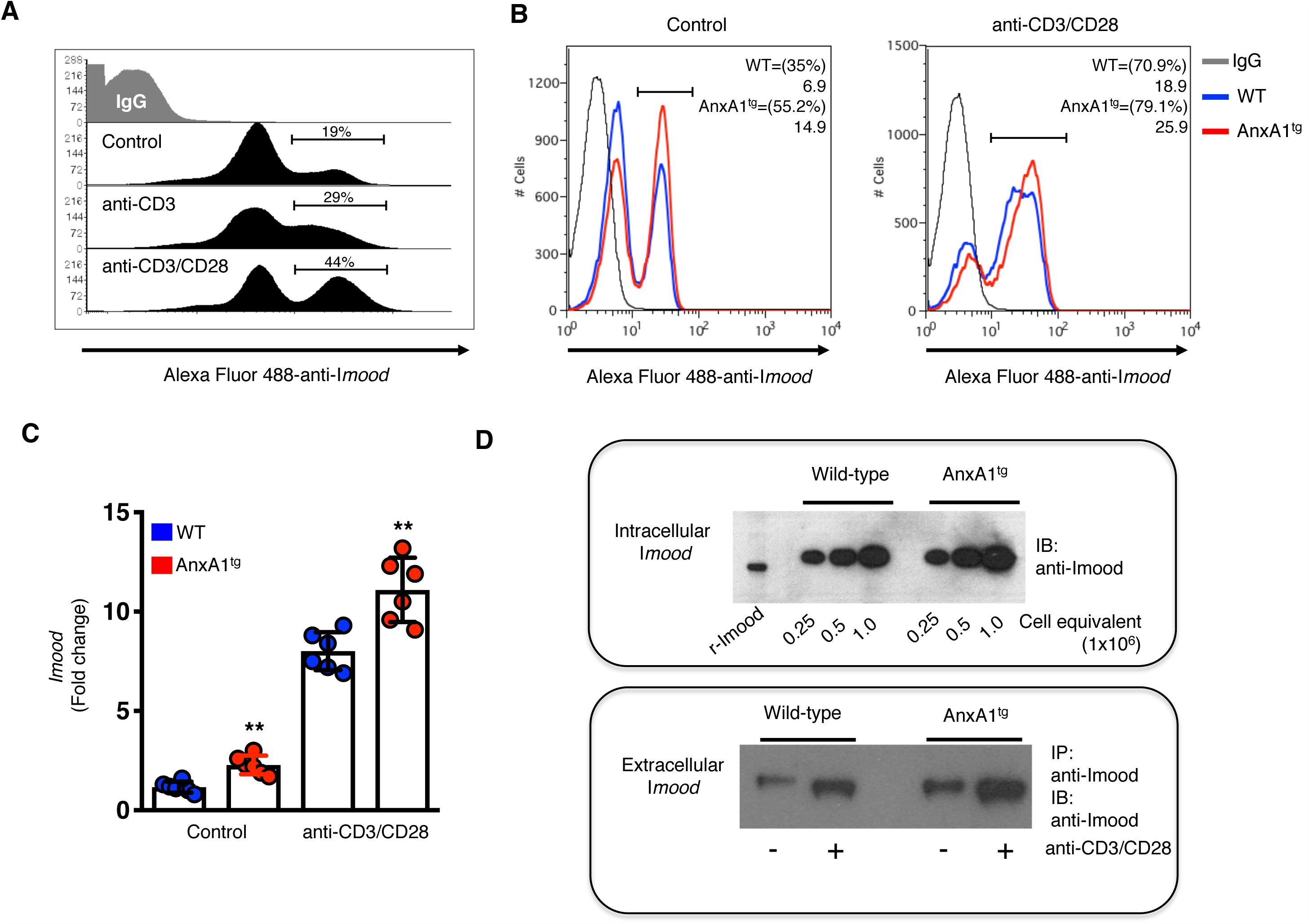
I*mood* expression in AnxA1^tg^ mice. (**A**) I*mood* intracellular staining of CD4^+^ T cells from C57BL/6 mice cultured overnight in complete medium (Control) or stimulated with 1μg/ml of plate-bound anti-CD3 or anti-CD3/CD28. The numbers in the gate represent the % of I*mood-*high expressing cells. The histograms show the results obtained with a single mouse and are representative or n=6-8 animals with similar results. (**B**) I*mood* intracellular staining of CD4^+^ T from wild type or AnxA1^tg^ mice cultured overnight in complete medium (Control) or stimulated with 1μg/ml of anti-CD3/CD28. The numbers in the plots show the % of I*mood-*high expressing cells (in the brackets) and the median fluorescence intensity of the gated region. The histograms show the results obtained with a single mouse and are representative or n=6 animals with similar results. (**C**) RT-PCR of I*mood* expression in CD4^+^ T cells from wild type and AnxA1^tg^ mice cultured overnight in complete medium (Control) or stimulated with 1μg/ml of plate-bound anti-CD3 or anti-CD3/CD28. Values are expressed as means ± SEM of n=6 mice. **p<0.01; ***p<0.001 indicate significant values compared to wild type control mice (Student’s *t*-test). (**D**) The top panel shows the western blotting of the whole cell lysates of indicated number of freshly isolated CD4^+^ T cells from wild type and AnxA1^tg^ mice. The bottom panel shows the levels of I*mood* immunoprecipitated from the cell culture medium of CD4^+^ T cells from wild type and AnxA1^tg^ mice cultured overnight in complete medium (Control) or stimulated with 1μg/ml of plate-bound anti-CD3 or anti-CD3/CD28. Membranes were immunoblotted with a polyclonal anti-I*mood* antibody and recombinant I*mood* (r-I*mood*) was used as control. The results shown are from a single mouse and are representative of six mice with similar results.

**Figure 6.**
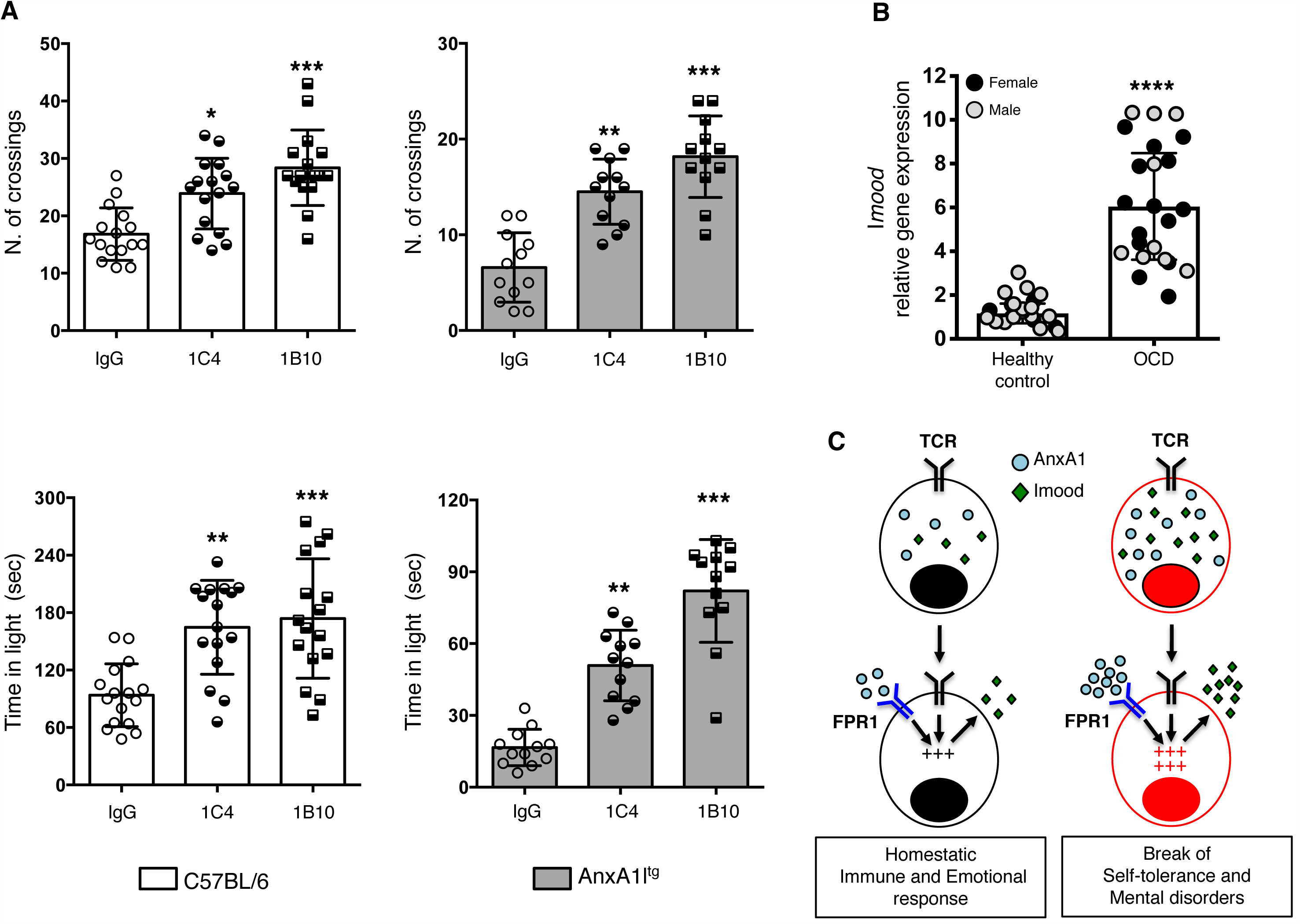
Anxiolytic effects of anti-Immuno-*mood*ulin antibodies. (**A**) AnxA1^tg^ (white bars) or C57BL/6 (grey bars) mice received an intraperitoneal injection (100ng/mouse) of anti-I*mood* antibodies 1B10 and 1C4 and then tested in the light and dark shuttle box test at day 7. The bar graphs show total number of crossings and the total time (seconds) spent in the lit area during a 5-minute trial. Values are expressed as means ± SEM of n=12 mice (AnxA1^tg^) or n=15 (C57BL/6) mice and are representative of three experiments with similar results. ****p<0.0001 indicate significant values compared with PBS-injected control mice (Mann–Whitney U-test). (**B**) Real time PCR analysis of I*mood* in peripheral blood mononuclear cells of patients diagnosed with OCD as described in Materials and Methods. Values are presented as individual data points ± SEM of 20 patients. Male and female subjects are coded as shown in the graph. ****p<0.0001 (Student’s *t*-test) (**C**) Schematic summery of the study showing how physiological levels of both AnxA1 and I*mood* play a homeostatic role regulating host immune response and emotional wellbeing. T cell activation causes the release of AnxA1 and the externalization of its receptor FPR. This signaling pathway integrates with the TCR and contributes to the regulation of the strength of TCR signaling and the level of T cell activation. Activated T cells express higher levels of I*mood*. The release of this protein by T cells contribute to a physiological state of lower mood that is similar to the sickness behavior observed following an infection. The increased level of expression of AnxA1 in T cells observed in patients suffering from autoimmune diseases is responsible for the lower threshold of T cell activation and the increased expression of I*mood*. The increased release of this protein in circulation leads to a state of higher anxiety and depression that is often observed in patients suffering from autoimmune conditions.

To explore the possible role of I*mood* in regulating anxiety behavior we administered the recombinant product of this gene (https://www.origene.com; Recombinant protein of human chromosome 8 open reading frame 42 [C8orf42], catalogue number TP313199) into wild type C57BL/6 mice. In all the experiments that followed, we used the light-dark shuttle box as a convenient screening test since this showed the highest difference between AnxA1^tg^ and wild type mice. As shown in Supplementary **Figure 5**, mice injected with 500ng of the intact (r-I*mood*) but not the denatured (95°C for 5min) protein (d-I*mood*) or PBS showed a significant increase in anxiety-like behavior. This change did not occur within hours of administration but started to be noticeable at day 3 post-injection (data not shown) and became highly significant at day 7. To confirm that I*mood* was indeed responsible for the increased anxiety-like behavior of AnxA1^tg^ mice, we administered a commercially available polyclonal antibody against this protein (https://www.novusbio.com/; catalogue number NBP1-93675) and control IgG to these mice. We also administered the same dose of antibodies and control IgG to wild type C57BL/6 to test its effect in control mice. As shown in **Supplementary Figure 6**, only AnxA1^tg^ mice that received the polyclonal anti-I*mood* antibodies but not those receiving the control IgG showed a significant increase of the time in the light (by ∼249%; p<0.001) and number of crossing (by ∼194%; p<0.001). Interestingly, administration of the same antibody in wild type C57BL/6 mice also caused a significant increase in time in the light (by ∼24%; p<0.05) and number of crossing (by ∼76%; p<0.01) suggesting there may be an endogenous I*mood* level which might regulate basal anxiety behavior of mice.

To further validate our hypothesis and increase specificity of action, we generated monoclonal antibodies against I*mood* using genetic immunization^48^ (**Supplementary Figure 7**). Congruent with previous data, administration of highly selective purified monoclonal anti-I*mood* antibodies 1B10 or 1C4 to AnxA1^tg^ significantly increased the number of crossing (by ∼123%, p<0.01 for 1C4; by ∼178%, p<0.001 for 1B10) and the time in the light (by ∼208%, p<0.01 for 1C4; by ∼396%, p<0.001 for 1B10) compared to IgG control. Thus, anti-I*mood* therapy rescues the phenotype of AnxA1^tg^ back to that of wild type mice.

Similar to what we observed with the polyclonal antibody, C57BL6 control mice administered with 1B10 and 1C4 showed significant increase in time in the light (by ∼75%, p<0.01 for 1C4; by ∼86%, p<0.001 for 1B10) and number of crossings (by ∼41%, p<0.05 for 1C4; by ∼68%, p<0.001 for 1B10). Collectively this data reveals I*mood* as an innovative target for therapeutic strategies to treat anxiety behavior in mice (**Figure 6A**).

### Increased expression of I*mood* in OCD patients

Finally, we searched for I*mood* expression in man. To this aim, we performed an initial retrospective screening of cDNA samples of PBMC obtained from obtained from 20 patients that have been diagnosed with Obsessive-Compulsive Disorders and 20 healthy controls. Both sociodemographic (gender, education, employment and marital status) and clinical (duration of illness, duration of untreated illness, family history of psychiatric disorders, psychiatric comorbidity and current treatments) variables have been reported in **Supplementary Table 3**. As shown in **Figure 6B**, I*mood* expression was significantly higher (approximately 6-fold) than controls in both male and female subjects.

## Discussion

A growing number of studies support the hypothesis that mood disorders can be driven by cellular and biochemical events that are rooted in the immune system ^10,49,50^. This evidence emerged from several experimental studies investigating how both the depletion of T cells in mice ^51,52^ or the repopulation of lymphopenic mice with T cells alter brain functions including cognitions ^53,54^, fear ^55^ and emotional states ^45,56^. Thus, in a study by Clark *et al.*, CD4^+^ T cells conferred anxiolytic and antidepressant-like effects in Rag-2^-/-^ mice ^57^. Using similar approaches, Scheinert *et al.* and Brachman *et al.* have shown that adoptive transfer of lymph node cells from chronically stressed mice were able to revert the anxious and depressive-like behavior of Rag-2^-/-^ mice ^58,59^.

The modulatory effect of T cells on behavior has also been explored using effector disease relevant T cells. Seminal studies by Beurel and Jope have shown that the number of effector Th17 cells was significantly higher in the brain of mice exhibiting depressive-like behavior or in mice subjected to chronic restraint stress ^60-62^. In line with the studies mentioned above, the same group has shown that adoptively transferred Th17 cells in Rag-2^-/-^ mice accumulated in the hippocampus of learned-helpless mice ^61^ and induced endogenous Th17 cell differentiation ^60^. This work has provided key evidence that autoimmune effector T cells play a fundamental role in regulating pathological processes other than autoimmunity. In the context of this study, these findings have laid the foundations for a further exploration of the possible mechanisms linking autoimmunity and mental disorders. In addition, the lack of novel therapeutic opportunities to treat mental health issues is very topical and we reason that detailed investigation of the mechanism(s) linking the immune system with behavioral responses could guide the development of new therapies.

We reasoned about the existence of novel mechanism and plausibly un-identified factors while noting the behavioral phenotype of mice which single anomaly was higher expression of AnxA1 in T cells. This transgenic tool was generated to further investigate the specific properties of this mediator on the adaptive immune response ^14,22,23^. In retrospective, our novel observations are aligned with the emerging notion that AnxA1 can regulate mental disorders. Genome-Wide Association Studies searches (GWAS Central at www.gwascentral.org) for AnxA1 reported about 62 studies on this protein and many of them on mental disorders. In fact, reports have shown a significant association between AnxA1 gene duplication and autism^63^ or single nucleotide polymorphism in AnxA1 gene and schizophrenia^64^, bipolar disorder or depression^65^. Most intriguingly, all these conditions have been often linked to immune dysfunctions or inflammatory conditions ^66-68^.

Moreover, AnxA1 is a ligand for the formyl peptide receptors (FPRs) ^69-71^. These prototype sensors of the innate immune system were initially identified as the cellular antenna for the capture of formylated peptides released by bacterial pathogens ^72-75^. Of interest, studies have shown that these receptors have more than one way to help the host sensing the danger. Their expression in the olfactory system allow mice to ‘sense’ the presence of infection-associated olfactory cues thus allowing animals to move away from the source of infection ^76-78^. Other lines of evidence support a role for FPR in regulating behaviour. Studies in knock out mice for both FPR1 and FPR2/ALX receptors have shown a significant reduction in anxiety ^79,80^ e.g. the opposite phenotype of AnxA1^tg^ mice. Putting everything together, FPR may represent a prototype of signalling molecules that influence the behaviour of the host at both cellular and physical levels with the ultimate goal of preserving it from the challenges of the external environments ^81^.

The anxiolytic effects of antibodies against I*mood* required days to be visible and statistically significant. A similarly delayed response was observed with recombinant I*mood* suggesting, in both cases, a downstream regulation of the expression of genes associated to anxiety rather than influencing directly the effects of neurotransmitters in the brain.

The time-lag effect for the emergence of modulatory effects on anxiety, with AnxA1^tg^ cells or following the administration of I*mood*, resonates well with the notion that classical drugs for the treatment of depression and anxiety present a delayed onset of 5 to 7 days for their clinical efficacy to be apparent ^82^. Recent studies have put forward the proposition that delayed onset could be linked to the time needed for the immune system to respond and/or adapt to the administration of these drugs^83^, reinforcing the hypothesis that the immune system regulates the emotional state *via* a homeostatic control of the expression of genes with direct effect on emotions.

This study opens a number of interesting and challenging questions that needs to be addressed to provide a full picture of how I*mood* might ultimately influence behavior. First of all, we would need to establish how I*mood* would enter the brain and what would be its receptor(s). Paradigm-shifting work by Kipnis and collaborators have very recently ascribed the meningeal lymphatic as “*a system capable of carrying fluid, immune cells, and macromolecules from the central nervous system*” ^84^. This system has been proposed as the vascular network that allows the circulation and exchange of soluble contents between the brain cerebrospinal fluid and interstitial fluid ^85,86^ and could be therefore the entry door for I*mood* in the CNS. Another interesting perspective would be the existence of an I*mood*-specific mechanics of transport present in epithelial cells of the choroid plexus and in endothelial cells of the blood-brain barrier like in the case of leptin ^87^. One surprising finding of our study is the existence of a small percentage of CD4^+^ T cells expressing high levels of I*mood* (Figure 5) and the increased number of these cells in the AnxA1^tg^ mice. It is tempting to speculate that these I*mood*-high cells could specifically recirculate through the brain and thus be directly involved in the regulation of brain gene expression profile. This would be particularly intriguing as we have observed a readily upregulation of this protein following TCR stimulation. The constitutive activation state of autoimmune T cells might justify, at least in part, the increased incidence of mental disorders in patients suffering these diseases. All these hypotheses could only be validated through systematic screening for I*mood* expression at both protein and mRNA levels in a large cohort of patients, with different types of mental disorders as well.

The possibility that I*mood* could represent a novel fine tuner of mental disorders would also offer the opportunity to have a novel biomarker of prognostic and diagnostic value. This would enable patient stratification for the correct mental disorder (e.g. those associated with an immune component) or identification of the right patient subgroup for specific drug treatment. As such, for those with higher expression of I*mood*, a combinatorial therapy with immunomodulators and I*mood* neutralizing antibodies like 1B10 might provide the unique opportunity to achieve a ‘healthy body in a healthy mind’. Along these lines, the identification of a protein mediator of emotional behavior and the availability of biological therapies that modulate its levels would represent a significant step forward for the treatment of mental disorders. Indeed, a biologic for the treatment of mental disorder would bypass several of the side effects associated with the daily administration of standard therapies mental disorders ^88-90^ as it would specifically act at the level of immune system rather than the CNS.

## Author contributions

NP generated the AnxA1^tg^ mice. GP and LR performed all the behavioral and immunological characterization of the AnxA1^tg^mice. SO and GB screened and tested antibodies against I*mood.* MO analyzed the microarray data. BD recruited OCD patients. CDA and FB assessed I*mood* expression in OCD patients. CM and MP contributed to the analysis and design of the study. FDA designed the study, analyzed the data, wrote the manuscript and contributed to all the stages of the experimental work.

## Data and materials availability

The data that support the findings of this study are available from the corresponding author upon reasonable request.

## Acknowledgments

We would like to thank Prof Kioussis for kindly providing us the CD2 cassette plasmid.

## Figures and Tables

**Supplementary Table 1. Differential gene expression profile whole brain from T cell specific AnxA1**^**tg**^ **mice.** The table shows a list of genes differentially modulated in AnxA1^tg^ brain compared to wild type controls. Probe ID is affymetrix ID. Fold changes (FC) is logged value.

**Supplementary Table 2. Differential gene expression profile of CD4**^**+**^ **T cells from T cell specific AnxA1**^**tg**^ **mice.** The table shows a list of genes differentially modulated in AnxA1^tg^ CD4^+^ T cells compared to wild type controls. Probe ID is Affymetrix ID. FC is logged value.

**Supplementary Table 3. Social and clinical variables of OCD subjects.**

**Supplementary Table 4. List of antibodies used for FACS.**

**Supplementary Figure 1. Generation of AnxA1**^**tg**^ **mice.** (**A**) Agarose gel electrophoresis of the AnxA1-Flag tagged construct cloned in two different backbone vectors (pCR2.1 and pcDNA3.1) and digested by *SmaI* to confirm the cloning of 1047 bp AnxA1 cDNA (indicated by the asterisk). (**B**) Schematic representation of the linearized construct that was used to generate T-cell specific AnxA1^tg^ mice. The construct was inserted in the *SmaI* site of the VACD2 vector – just between the CD2 promoter sequence and CD2 locus control region (LCR) cassette. (**C**) The two agarose gels show the results of the PCR screening (for further details see Supplementary Material and Methods) of the 22 pups obtained from the pronuclear injection of VACD2-AnxA1-Flag construct. The top gels show the 3 mice that resulted positive for the transgene while the bottom show the PCR screening for the endogenous AnxA1 as control. (**D**) Western blotting of the whole cell lysate from purified equal number (1×10^6^) of T cells of wild type and AnxA1^tg^ mice showing increased expression of AnxA1 protein in transgenic mice.

**Supplementary Figure 2. Immune repertoire of AnxA1**^**tg**^ **mice.** (**A**) Representative dot plots showing CD4/CD8 staining of thymus, spleen and lymph node T cells of wild type and AnxA1^tg^ T cells. (**B**) The bar graphs show the total cell number of cells purified from wild type and AnxA1^tg^ mice as described in Supplementary Materials and Methods, section “Flow Cytometric analysis”. Values are expressed as means ± SEM of n=6 mice for each group and are representative of four experiments with similar results. ****p<0.01 (Student’s *t*-test) indicates significant values compared to wild type control mice.

**Supplementary Figure 3. Autoimmune-prone phenotype and heightened anxiety-like behavior AnxA1**^**tg**^ **mice.** (**A**) Control and AnxA1^tg^ mice derived from the second founder (red line; see Results section for more details) were immunized with MOG_35-55_ and CFA and monitored daily for clinical signs of EAE (top left panel) or weight gain/loss (top right panel) for 23 days. Results show means ± SEM from n=10 mice per group and are representative of three separate experiments with similar results. **** p<0.001, (two-way ANOVA followed by Bonferroni multiple-comparison test) indicates significant values compared to wild type control mice. (**B**) The bar graphs on the left show the total duration (seconds) of digging and total number of buried marbles, during a 10-minute trial of the marble burying test. Those in the middle show the total time (seconds) spent in the lit area and total number of crossing during a 5-minute trial in the light/dark shuttle box test. The bar on the right show total number of squares crossed and the latency (seconds) to the first rearing during a 5-minute session in the open field test. Values are expressed as means ± SEM of n=15 mice and are representative of three separate experiments with similar results. ***p<0.001; ****p<0.0001 indicate significant values compared to wild type control mice (Mann–Whitney U-test).

**Supplementary Figure 4. Increased signs of organ inflammation in AnxA1**^**tg**^ **subjected to pristane-induced lupus.** Control and AnxA1^tg^ mice received an intraperitoneal injection of pristane to induce a lupus-like disease. After 4 weeks mice were culled and dissected to collect spleens and lungs. Tissues were fixed with 4% PFA and then stained with hematoxylin and eosin. The panels in **A** show a photo picture of the spleen (left) and it relative histological section (right). The panels in **B** show a photo picture of the lung (left) and it relative histological sections (middle and right panels, 4x and 20x magnifications). Results are from a single mouse and are representative of n=10 mice with similar results.

**Supplementary Figure 5**. C57BL/6 mice were injected with PBS or r-I*mood* or denaturated r-I*mood* (d-I*mood)* (500ng, i.p.; https://www.origene.com; Recombinant protein of human chromosome 8 open reading frame 42 (C8orf42), catalogue number TP313199) and tested at day 7 post injection. The bar graphs show the total time (seconds) spent in the lit area and total number of crossings during a 5-minute trial. Values are expressed as means ± SEM of n=12 mice and are representative of three experiments with similar results. ****p<0.01 indicates significant values compared to PBS-injected control mice. §§p<0.01; §§§p<0.001 indicate significant values compared to d-I*mood*-injected mice. (Mann–Whitney U-test).

**Supplementary Figure 6. Effect of anti-I*mood* polyclonal antibody on anxiety-like behavior of C57BL/6 and AnxA1**^**tg**^ **mice.** C57BL/6 and AnxA1^tg^ mice received an i.p. injection of polyclonal anti-I*mood* or IgG control antibodies (500ng i.p.) and then tested at day 7. The bar graphs show the total time (seconds) spent in the lit area and total number of crossings during a 5-minute trial. Values are expressed as means ± SEM of n=12 mice and are representative of three experiments with similar results. ***p<0.001; ****p<0.0001, indicate significant values compared to IgG-injected control mice (Mann–Whitney U-test).

**Supplementary Figure 7. Screening and identification of 1B10 and 1C4 monoclonal anti-I*mood* antibodies.** (**A**) Aliquots (50ng) of recombinant I*mood* were loaded on an SDS-page gel and then transferred on PVDF membranes as detailed in Materials and Methods. Membranes were immunoblotted with the supernatants from different hybridoma cultures (code names indicated on the top of the top panel). Thereafter, the same membranes were stripped and immunoblotted with a commercially available anti-I*mood* antibody (bottom panel). (**B)** Aliquots (50ng) of recombinant I*mood* were immunoprecipitated with hybridoma supernatants and then loaded on an SDS-page gel as detailed in Materials and Methods. Membranes were immunoblotted with a commercially available anti-I*mood* antibody.

## SUPPLEMENTARY MATERIALS

### Reagents

Unless otherwise stated, all the chemicals were purchased from Sigma. For the convenience of the readers all the antibodies used in the study have been reported in Table 4 as well as in the figure legends.

### Animals and husbandry

Wild type C57BL/6J (B6-CD45.2) and B6.SJL-*Ptprca Pepcb/BoyJ* (B6-CD45.1) were purchased from Charles River. AnxA1^tg^ mice were generated in the in-house transgenic mouse facility of Queen Mary University of London by pronuclear injection.

Mice were housed in groups of 6 per cage under specific-pathogen-free conditions and with free access to food and water. Mice were housed for at least 7 days prior to testing. All experiments were performed during the light phase of the light-dark cycle and no more than 2 tests per day were performed. The results presented in this manuscript were obtained using male mice (6-8 weeks old) because - unlike in humans - both the intensity and incidence of inflammatory/autoimmune disease are more pronounced in male mice. However, the difference we observed in terms of both immune response and behavioral changes were confirmed and reproducible in female mice (data not shown). All tests were conducted under license from the Home Office and according to the UK Animals (Scientific Procedures) Act, 1986. All experiments were approved and performed under the guidelines of the Ethical Committee for the Use of Animals, Bart’s and The London School of Medicine and Home Office Regulations (57) (PPL 80/8714).

To monitory the transgenic colony, genomic DNA were extracted from ear clips by using REDExtract N-AMP –XNAT kit (Sigma, UK) and analysed by PCR with the following specific primers for AnxA1^tg^: forward primer 5’-GTATGGAATCTCTCTTTGCCAAGC-3’; reverse primer is 5’-ACHGATATGCACATCAGGAGGG-3’ (Thermo Scientific, UK). The parameters of the PCR reactions are: initial denaturation at 94°C for 3min followed by 30 cycles of denaturation at 94°C for 45sec, annealing at 60°C for45sec and extension time at 72°C for 15sec, and afterwards a final extension step at 72°C for 7 min.

All the behavioral tests were video-recorded and analyzed double-blind during the light phase of the light-dark cycle, as previously described^91^. The scoring and analysis of the data was hand-made by at least 2 blinded independent experimenters. All the efforts were made to minimize mouse discomfort in these behavioral experiments. Mice were brought to the testing room at least 30 minutes before the start of the test session to allow habituation to the testing environment. All behavioral equipment was thoroughly cleaned and sanitized at the end of the day and in between tests. Unless otherwise specified, standard lighting (about 50 lux) and quiet conditions were maintained throughout each experiment.

### Flow cytometric analysis

Single cell suspension of thymocytes or lymphocytes from spleen and lymph nodes were obtained as previously described ^19,45^. Briefly, tissues were gently desegregated in RPMI medium supplemented with 100U/ml of penicillin and streptomycin (PAA laboratories, Buckinghamshire, UK) using a 70μm mesh cell strainer (Falcon, UK) and the piston of a 5ml syringe. Cell suspensions from spleen and lymph nodes were layered onto Histopaque-1077 (Sigma-Aldrich, Dorset, England) in a ratio 3:1 and centrifuged at 400g for 10 mins to isolate mononuclear cells. The resulting buffy coat was collected and washed twice with RPMI supplemented with 100U/ml of penicillin and streptomycin (PAA laboratories, Buckinghamshire, UK) and then suspended in FACS buffer (phosphate-buffered saline containing 5% fetal calf serum and 0.02% of NaN_2_) for further analysis. Thymocytes were washed twice with RPMI supplemented with 100U/ml of penicillin and streptomycin (PAA laboratories, Buckinghamshire, UK) and then suspended in FACS buffer. Aliquots of cell suspensions were used to evaluate the total number of cells using a standard hematocytometer. Cells were stained in 100 µl of FACS buffer containing the following fluorochrome-conjugated antibodies: anti-CD3 (clone 145-2C11), anti-CD4 (clone GK 1.5), anti-CD8 (clone 53-6.7) (all from eBioscience, San Diego, CA, USA). Cells were labeled with the appropriate concentration of conjugated antibodies for 1 h at 4°C. After labeling, cells were washed and analyzed using FACScalibur flow cytometer. Results were analyzed using the FlowJoTM software (Tree Star, Ashland, OR, USA, Oregon Corporation).

### T cell activation assay

Lymph node T cells (1×10^5^ cells/200 µl) were incubated with medium alone or stimulated by plate-bound anti-CD3 (clone 145-2C11; eBioscience) and anti-CD28 (clone 37.51; eBioscience) for 24 hours in 96-well plates. For CD25 and CD69 upregulation, lymph node T cells were stimulated with plate-bound anti-CD3 and anti-CD28 as indicated in the figure. After 16 hours, the cells were stained with PE-conjugated anti-CD69 (clone H1.2F3) and FITC-conjugated anti-CD25 (clone PC61.5) diluted in FACS buffer (PBS containing 1% FCS and 0.02% NaN_2_). The polyclonal antibody against Tdrp (I*mood*) was purchased from Novusbio (catalogue number NBP1-93675). The secondary antibody was a goat anti-rabbit Alexa Fluor 488-conjugated IgG from Abcam (catalogue number ab150077). Intact cells were gated by using forward and side scatter and analyzed with the FlowJoTM software (Tree Star, Ashland, OR, USA, Oregon Corporation). IL-2 production was measured after 24 or 48 hr of stimulation using a standard ELISA kit and according to the manufacturer’s instructions (eBioscience).

### Intracellular staining and cytometric bead assay

Th cell phenotype was studied by intracellular staining. Lymphocytes were isolated from MOG_35-55_ immunized mice from peripheral lymphoid organs and spinal cord. Lymphocytes (10×10^6^ cells/ml) from lymph nodes and spleen were cultured for 72 hours with either medium alone (CTRL) or with anti-CD3 (clone 145-2C11; eBioscience) and anti-CD28 (clone 37.51; eBioscience) antibodies (1µg/ml; plate bound) or with the specific antigen MOG_35-55_ (100µg/ml). At third day, the cells were pelleted and the supernatants stored at - 20°C. Subsequently, the cells were re-challenged with concanavalin A (ConA, 5µg/ml; Sigma) in presence of protein transport inhibitor Brefeldin A (1:1000; eBioscience) for 4 hours. Mononuclear cells isolated from the spinal cords instead were directly triggered with ConA and Brefeldin A after collection.

Cells were pelleted and then stained for CD4 (clone GK 1.5; eBioscience) for half an hour and fixed with 1% PFA for 10 minutes. Thereafter, cells were permeabilized and stained for 30 min in permeabilization buffer (eBioscience) containing conjugated antibodies (all from eBioscience) for cytokines (dil: 1:250) such as IFNγ (clone XMG1.2), IL-17 (eBioTC11-18H10.1), GM-CSF (clone MP1-22E9) and IL-10 (clone JES5-16E3). Finally, cells were washed and suspended in FACS buffer for flow cytometer analysis.

### Cytometric bead array

Cytokine production was measured by bead-based analytic assay in flow cytometry. We used a custom-designed Mouse Th1/Th2/ Th17 /Th22 13plex kits supplemented with antibody-bounded beads for GM-SCF and IL-23 (eBioscience). Each sample (25µl of cell culture supernatant) was incubated with 50µl bead mixture and 50µl mix of antibodies conjugated with biotin for 2h. After two washes, streptavidin PE conjugated antibodies was added and samples were let rocking for 1 hour in dark. Finally, samples were washed and stored overnight at 4°C. Standards diluted serially for 7 times were prepared and processed at the same time. Samples were analysed using a BD LSR Fortessa using the following voltage settings: FSC 444, SSC 250, YG-585/15 channel 430, R-670/14 channel 400.

### Histology

Intact spinal cords were first fixed in 4% paraformaldehyde for 72 h and then incubated with decalcifying solution containing EDTA (0.1 mM in PBS) for 14 days prior to paraffin embedding. Tissues sections (5 µm) were deparaffinized with xylene and stained with hematoxylin and eosin (H&E) by our in-house histology facility (https://www.bci.qmul.ac.uk/en/research/lab-facilities/pathology). Photos of the sections were acquired using an Olympus BX61 available at the in-house pathology facility. In all cases, a minimum of three sections per animal was evaluated. Phase-contrast digital images were taken using the Image Image-Pro (Media Cybernetics, Rockville, MD, USA) analysis software package.

### MOG_35-55_-induced Experimental Autoimmune Encephalomyelitis

Male C57BL/6 mice received an intradermal injection of 300 μg of MOG_35-55_ (MEVGWYRSPFSRVVHLYRNGK, synthetized by Cambridge Research Biochemicals, UK; Cambridge Research Biochemicals) emulsified in Complete Freund’s adjuvant (CFA; Sigma-Aldrich, Dorset, England) and two doses of 500ng of pertussis toxin (PTX; Sigma-Aldrich, Dorset, England) at day zero and day two as previously reported ^20^. The severity of the disease was scored on a scale of 0 to 6 with 0= no neurological signs, 1=tail weakness, 2= tail paralysis, 3= loss of righting reflex (the mouse can no longer right themselves after being laid on their back), 4= hind leg paralysis, 5= quadriplegia and 6= death.

### Leukocytes isolation from central nervous system

Vertebral columns were dissected from the lumbar to the cervical region and washed several times in PBS to remove blood trace. Spinal cords were extracted by hydro pressure in the spinal canal by using a 2ml syringe and 19-gauge needle. Subsequently, tissues were torn apart in sterile PBS by mechanical pressure through a 70µm mesh cell strainer (Falcon). Mononuclear cells and lymphocytes were isolated by density gradient centrifugation in Percoll (GE Healthcare). In detail, cells were pelleted at 400xg for 5 min and suspended in a 30% Percoll solution. The 30% Percoll solution was carefully layered onto a 70% Percoll solution in a ratio 1:2 and centrifuged at 500xg for 30 min. In this density gradient mononuclear cells sediment at the interface between 30% and 70% Percoll layers. About 2-3ml of interface solution was collected only after the fatty layer at the top of the centrifuge tube was carefully removed. The purified mononuclear cells were washed twice in RPMI supplemented with 100U/ml of penicillin and streptomycin and 10% of FCS (Invitrogen).

### Pristane-induced lupus

Wild type or AnxA1^tg^ male mice received received a single 0.5 ml i.p. injection of sterile pristane (Sigma-Aldrich, St Louis, MO, USA). Weight was recorded every other day or every 3 days for over 5 weeks. In some experiments, mice were culled after 4 weeks and the spleen and lung collected for histological analysis. The weight of the spleen was also measured to compare the level of splenomegaly between wild type or AnxA1^tg^.

### Digging and marble-burying tests

Marble-burying and digging tests were carried out as described previously ^80^ with some modifications. Briefly, mice were individually placed in a clear plastic box (48cm × 39cm × 31cm) filled with approximately 5-cm-deep wood chip bedding lightly pressed to give a flat surface. The same bedding substrate was used for all the mice and flattened after each test. Twenty glass marbles were placed on the surface in five rows of 4 marbles each. The latency to start digging, the number of digging bouts and the number of buried marbles (to 2/3 their depth) were recorded during the 10 min test.

### Climbing activity test

The climbing test is used to assess vertical activity and exploratory behavior. The test was performed as previously described ^80^. Briefly, mice were placed, one at a time, on a thin layer of fresh wood chip bedding on a laboratory bench and covered with a cylindrical climbing mesh (60 cm × 30 cm base diameter). They were each observed and recorded for 5 minutes. The number of climbing events and total duration of climbing activity was assessed. The criterion for climbing was for a mouse to have all 4 feet on the wire mesh while a climb terminated as soon as one foot touched the bench. This test was conducted in the late afternoon, when mice are known to be more active.

### Light-dark shuttle box

In this test exploratory activity reflects the combination of hazard and risk avoidance ^41^. The apparatus consisted of a 45cm × 20 cm × 21 cm box, divided into two distinct compartments: one third (15 cm long) painted black, with a black lid on top, the remaining two thirds painted white and uncovered. A 2.5 cm × 2.5 cm opening joined the two compartments. One side of the bright box was transparent to enable behavioral assessment and the averseness of this compartment was increased by additional illumination supplied by a 50 W lamp placed 45 cm above the centre of the box floor. The test was performed in accordance with a previous published protocol ^41^. Each mouse was placed in the bright compartment, facing away from the opening and allowed to explore the box for 5 minutes. Dependent variables included the time spent in the light area, latency to cross to the dark area (all four paws in) and the total number of transitions between compartments. The apparatus was cleaned after each trial.

### Open field activity test

The open filed test was performed as described previously ^80^. The open field consisted of a white PVC arena (50cm×30cm) divided into 10cm×10cm squares. Mice were brought into the experimental room 15min before testing. Each mouse was placed in one of the corner squares facing the wall, observed and recorded for 5min. The total number of squares crossed, latency to the first rear and the total number of rears were recorded. After each test, the arena was cleaned with water to attenuate and homogenize olfactory traces.

### Real-time polymerase chain reaction

Total RNA was extracted from brains of wild type and AnxA1^tg^ mice RNeasy Microarray Tissue Mini Kit (Qiagen, West Sussex, UK) while for the purified CD4^+^ T cells we used RNeasy Mini Kit from the same manufacturer. Meninges were removed by gently rolling the brains on Whatman paper. Total RNA was reverse transcribed using 2 mg oligo(dT)15 primer, 10 U AMV reverse transcriptase, 40U RNase inhibitor (all from Promega Corporation, Madison, WI, USA) and 1.25mM each dNTP (Bioline, London, UK) for 45 min at 42°C. Real-time polymerase chain reaction was carried out by using ABsoluteTM QPCR ROX Mix (Thermo Scientific, Epsom, UK) and fluorescent QuantiTect primers (Qiagen, West Sussex, UK). The human and murine primers for Tdrp were HsTDRP1SG and MmTdrp1SG. Cycling conditions were set according to the manufacturer’s instructions. Sequence-specific fluorescent signal was detected by 7900HT Fast Real-Time PCR System (Applied Biosystems, Warrington, Cheshire, UK). mRNA data were normalized relative to glyceraldehyde 3-phosphate dehydrogenase and then used to calculate expression levels. We used the comparative Ct method to measure the gene transcription in samples. The results are expressed as relative units based on calculation of 2^-ΔΔCt^, which gives the relative amount of gene normalized to endogenous control (glyceraldehyde 3-phosphate dehydrogenase) and to the sample with the lowest expression set as 1.

Total RNA was also isolated from PBMCs of OCD subjects and healthy controls according to the method of Chomczynski and Sacchi ^92^. RT-PCR reactions were performed using the RevertAid H Minus First Strand cDNA Synthesis Kit (Thermo Scientific, Waltham, MA, USA). The relative abundance was assessed by RT-qPCR using iQ SYBR Green Supermix (Hercules, CA, USA) on a DNA Engine Opticon 2 Continuous Fluorescence Detection System (MJ Research, Waltham, MA, USA). To provide precise quantification of the initial target in each PCR reaction, the amplification plot was examined and the point of early log phase of product accumulation defined by assigning a fluorescence threshold above background, defined as the threshold cycle number or Ct. Differences in threshold cycle number were used to quantify the relative amount of the PCR targets contained within each tube. After PCR, a dissociation curve (melting curve) was constructed in the range of 60 to 95 °C to evaluate the specificity of the amplification products. The relative expression of different amplicons was calculated by the delta-delta Ct (ΔΔCt) method and converted to relative expression ratio (2^−ΔΔCt^) for statistical analysis. All human data were normalized to the endogenous reference genes β-actin and GAPDH combined.

### Microarray analysis

Total RNA was hybridized to Affymetrix Mouse Gene 1.0 ST array chips at UCL Genomics (London, UK) with standard Affymetrix protocols, using GeneChip Fluidics Station 450, and scanned using the Affymetrix GeneChip Scanner (Affymetrix, Santa Clara, CA, USA). Data was normalized by robust multiarray average (RMA) of the Bioconductor package, affy. Relevant genes were filtered by excluding those without an Entrez ID and those with low expression levels less than 100 by non-logged value. T-statistics were applied across the data set using the Bioconductor package Limma, and differentially expressed genes were identified by P < 0.05 (non-adjusted P value).

### Western blotting analysis

Lymph node and splenic T cells or purified CD4^+^ T cells were stimulated as indicated in figures. To obtain the whole cell lysate, cells were resuspended in ice-cold lysis buffer (1% NP-40, 20 mM Tris pH 7.5, 150 mM NaCl, 1 mM MgCl_2_, 1 mM EGTA, 0.5 mM PMSF, 1 µM aprotinin, 1 µM leupeptin, 1 µM pepstatin, 50 μM NaF, 1 μM NaVO_4_, 1 μM µ-glycerophosphate) and left on ice for 5 mins. Cell lysates were centrifuged at 13,000 rpm at 4°C for 5 mins and the post-centrifugation supernatants collected, mixed with 6×Laemmli buffer (Invitrogen) and stored at −20°C.

To immunoprecipitate extracellular released I*mood*, 5 µl of rabbit polyclonal antibody against Tdrp (Novusbio; catalogue number NBP1-93675) and 35 µl of protein A/G sepharose beads (Santa Cruz) were added to 500 µl of culture supernatants obtained from 1×10^7^/ml CD4^+^ T cells stimulated with plate-bound anti-CD3/CD28 (1µg/ml) for 24 hrs. Samples were incubated overnight at 4°C under continuous rotation and then washed with cold PBS for 3 times. Soon after the washes, samples were denatured with 35ml of 6×Laemmli buffer (Invitrogen).

Lysates and immunoprecipitates were subjected to electrophoresis on SDS-12% polyacrylamide gel (Novagen). After subsequent transfer onto polyvinylidene difluoride Immobilon-P Transfer membranes (Millipore, Watford, UK), these were incubated overnight with antibodies diluted in Tris-buffered saline solution containing Tween-20 (TTBS: 130 mM NaCl; 2.68 mM KCl; 19 mM Tris-HCl; 0.001% v/v Tween-20, pH 7.4) with 5% nonfat dry milk (Marvel) at 4°C. Immunoblotting and visualization of proteins by enhanced chemiluminescence (ECL; Amersham Biosciences, Piscataway, NJ, USA) were performed according to manufacturer’s instructions. The recombinant Tdrp used as positive control was purchased from Origene (https://www.origene.com; Recombinant protein of human chromosome 8 open reading frame 42 (C8orf42), catalogue number TP313199).

### Genetic Immunization

Genetic immunization is an antibody generation platform offered as service by Aldveron (http://www.aldevron.com). This technology is most suitable to generate neutralising antibodies because native proteins are expressed *in vivo* with normal posttranscriptional modifications. The target is cloned into one of Aldveron proprietary immunization vectors and introduced into the host organism for gene expression. To help identify positive antisera and hybridomas, screening without purified antigen is made possible with the use of Aldveron proprietary screening vectors.

The procedure followed for the generation of 1B10 and 1C4 is herein briefly described. Six rats were immunised with the immunisation vector pB8-TDRP-mur containing TDRP cDNA sequence. The immune serum was taken at day 24 of the immunisation protocol, after 4 genetic applications. Sera, diluted in PBS 3% FBS, were tested by flow cytometry using mammalian cells transiently transfected with the murine Tdrp cDNA cloned into an Aldevron proprietary expression vector (pB1-TDRP-mur). A goat anti-rat IgG R-phycoerythrin conjugate (Southern Biotech, #3030-09) at 10 µg/ml was used as a secondary antibody. As negative control, mammalian cells transfected with an irrelevant cDNA cloned in the same expression vector were used. Cell surface expression of the screening vector pB1-TDRP-mur was analysed by flow cytometry using an anti-tag antibody and a goat anti-mouse IgG R-phycoerythrin conjugate (Southern Biotech, #1030-09) at 10 µg/ml as a secondary antibody.

### Statistics

According to the nature of the data obtained, a Student’s *t* test (2-tailed), or a 1 or 2-way analysis of variance (ANOVA) was performed. Time-course observations were analysed with two-way ANOVA followed by Bonferroni multiple-comparison test. Behavioural data was analysed *via* nonparametric analysis using the Mann-Whitney U test. All statistical analysis was performed using GraphPad PRISM software v8.0, with the exception of microarray analysis which was carried out as stated in 2.13.3 using the software package LIMMA (Bioconductor). Data was analysed for normality using the D’Agostino-Pearson omnibus normality test.

