## Supplemental data for "Immuno-*moodulin*: a new anxiogenic factor produced by autoimmune-prone T cells"

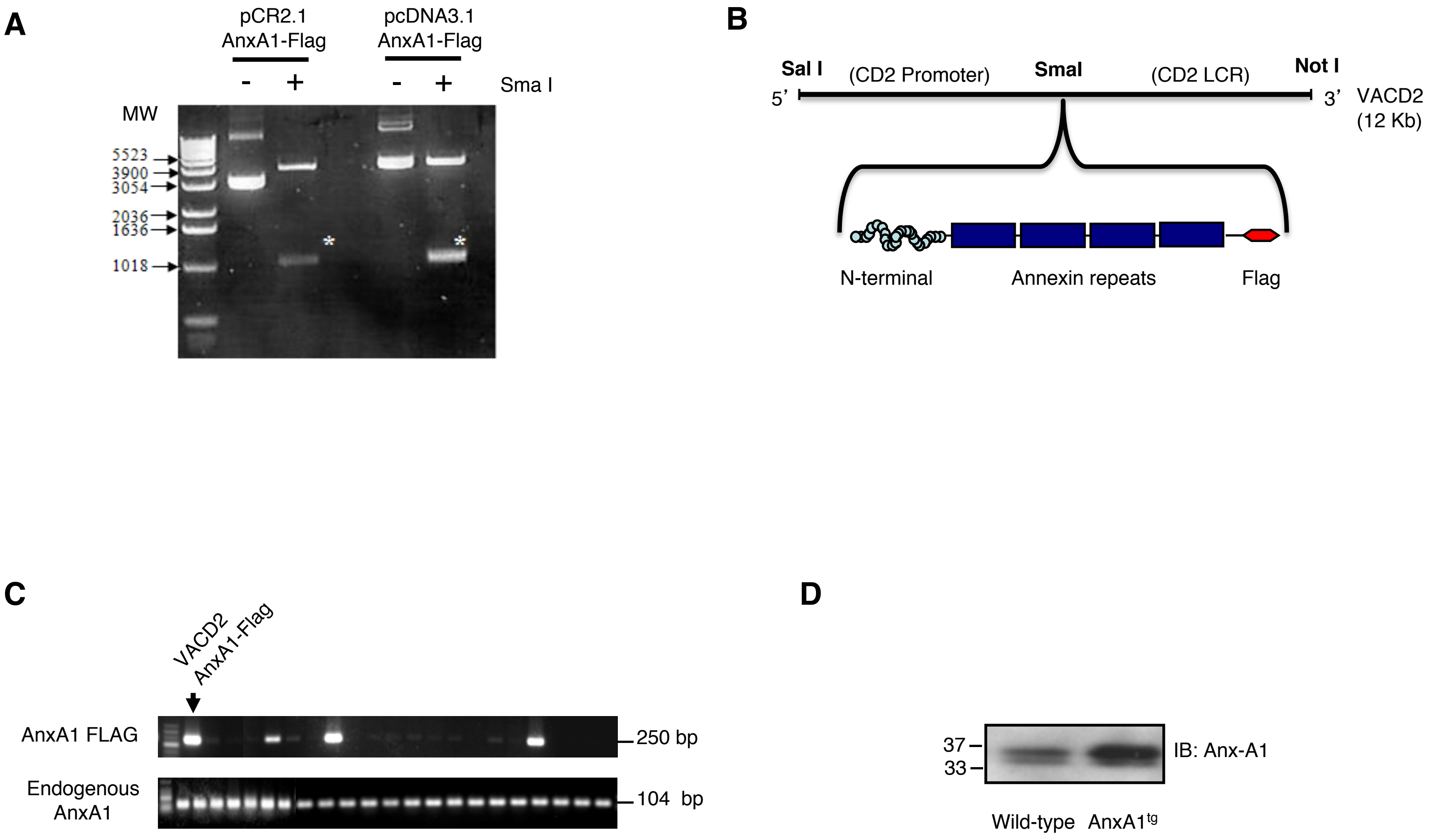

Piras *et al.*, Supplementary Figure 1

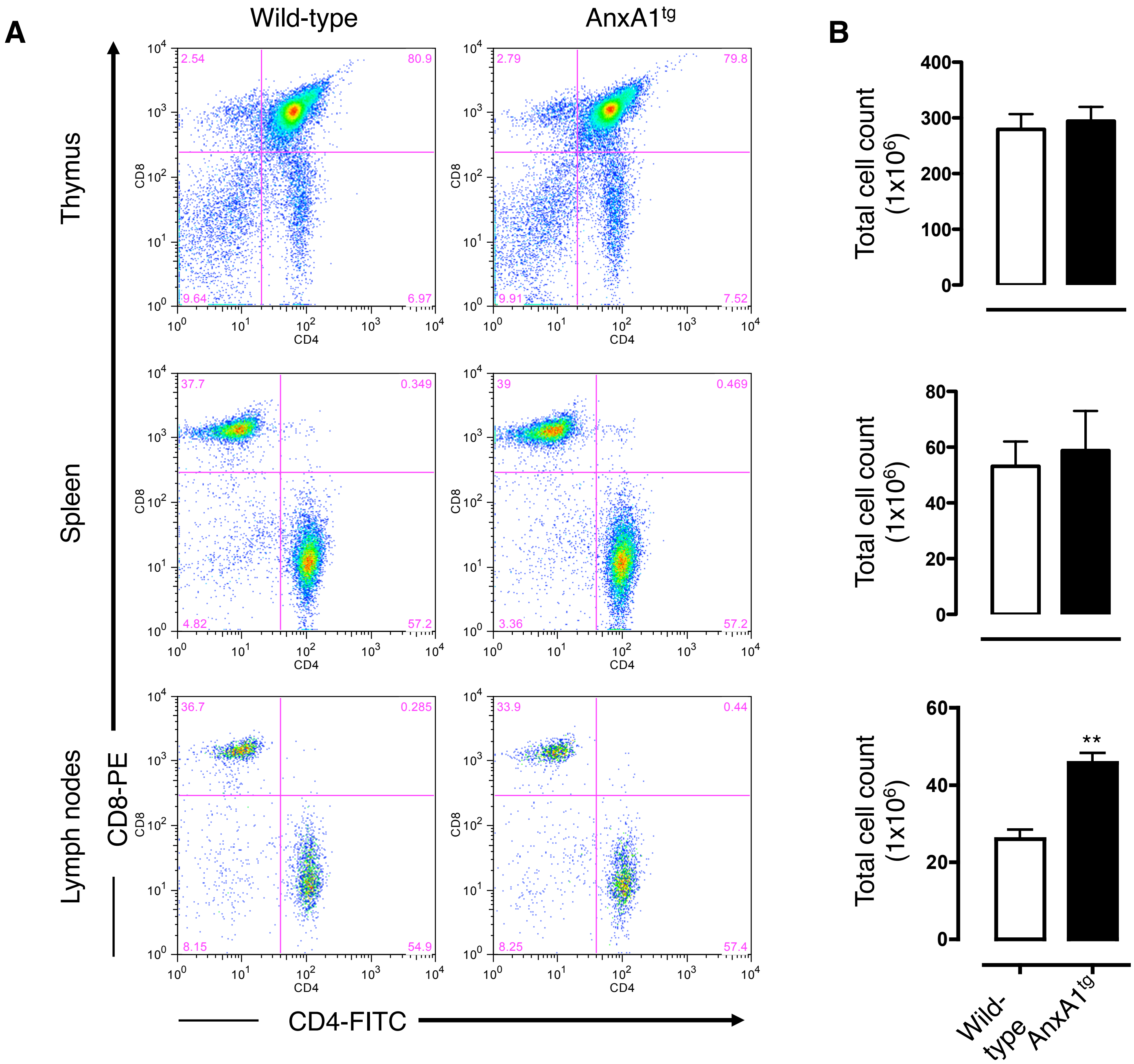

Piras *et al.*, Supplementary Figure 2

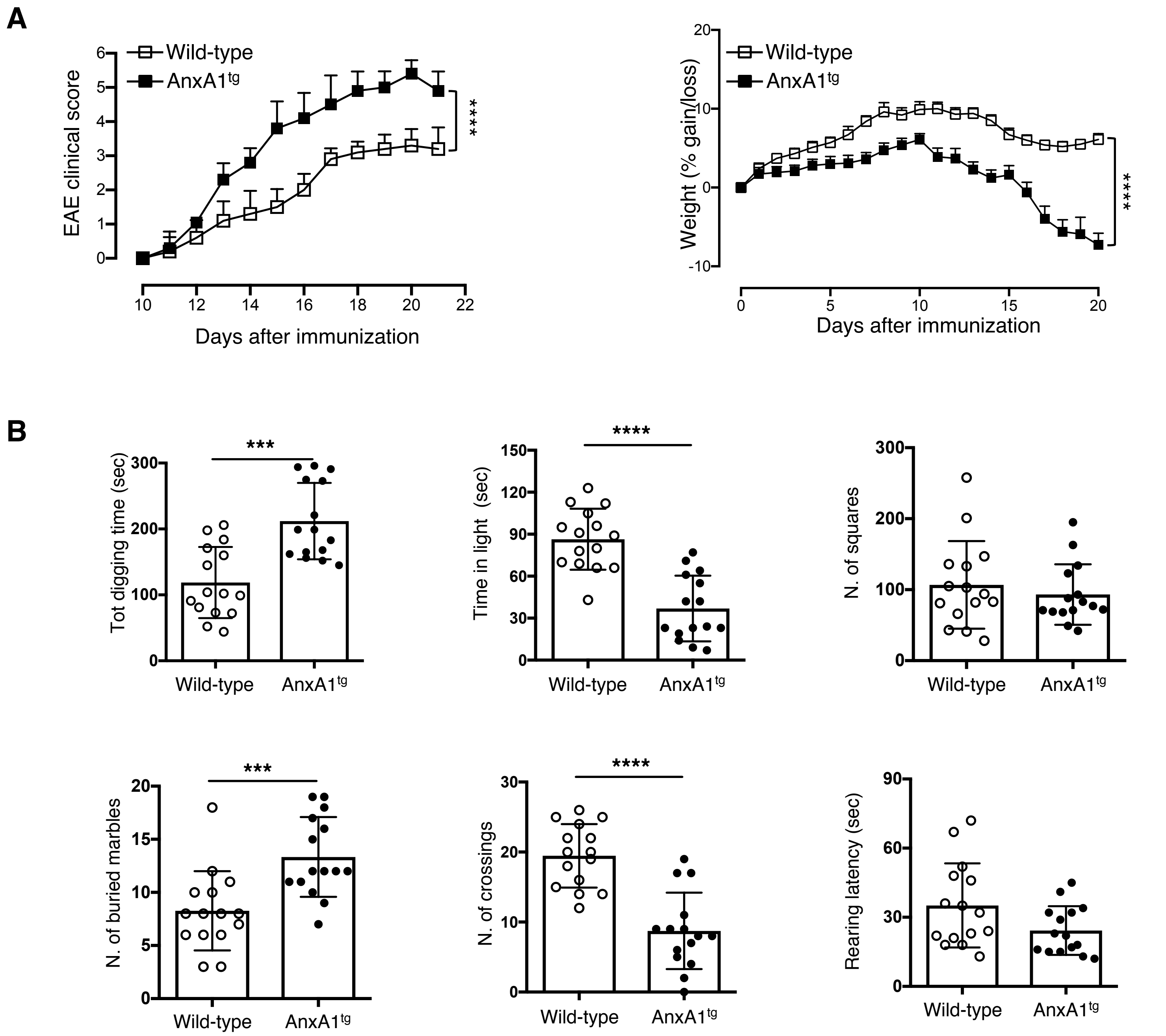

Piras *et al.*, Supplementary Figure 3

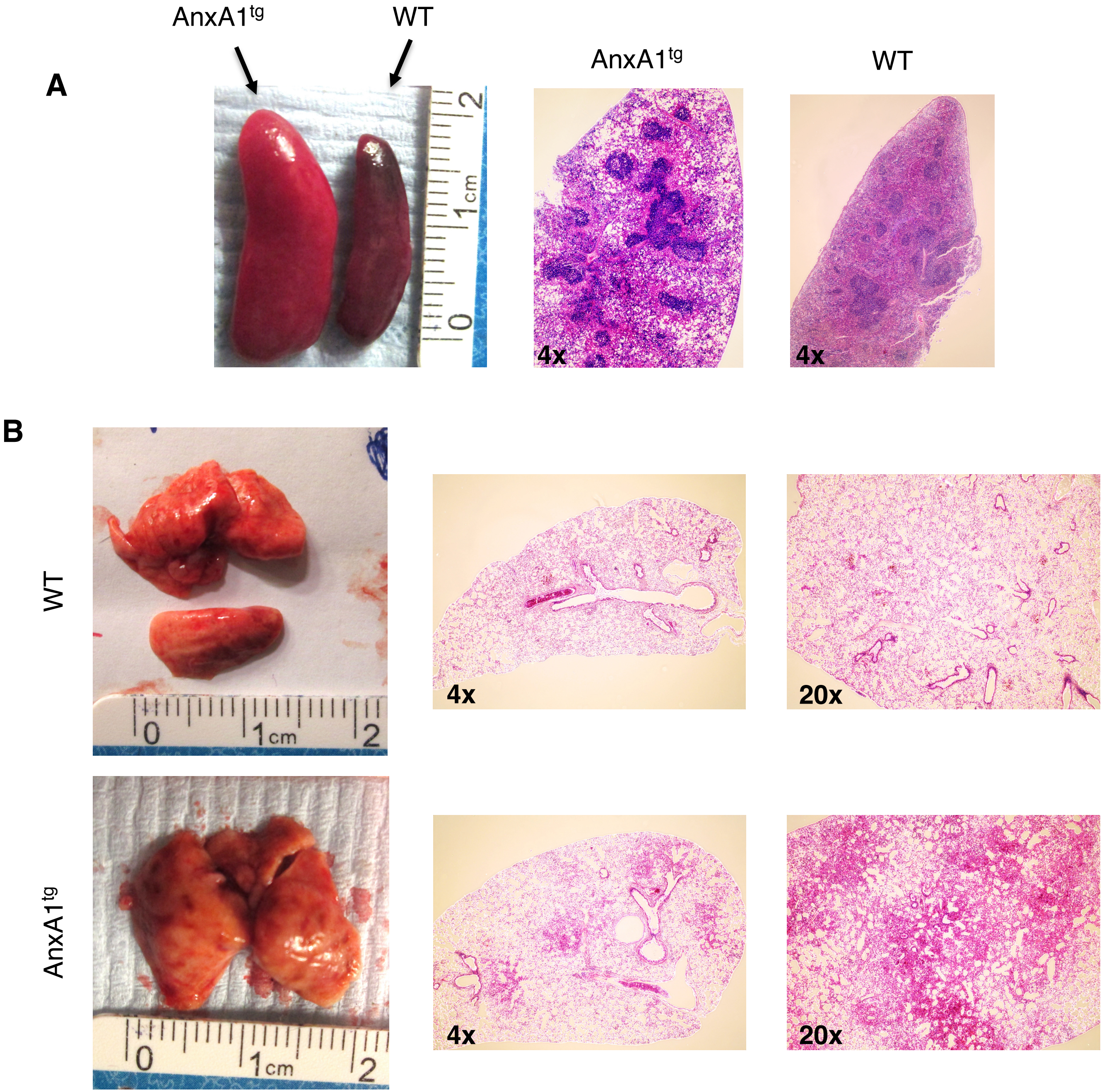

Piras *et al.*, Supplementary Figure 4

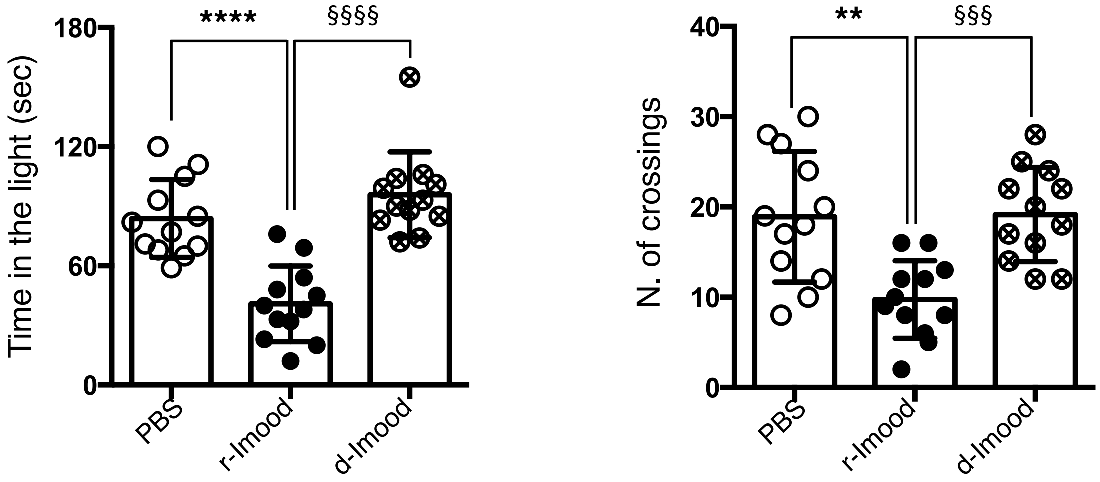

Piras et al., Supplementary Figure 5

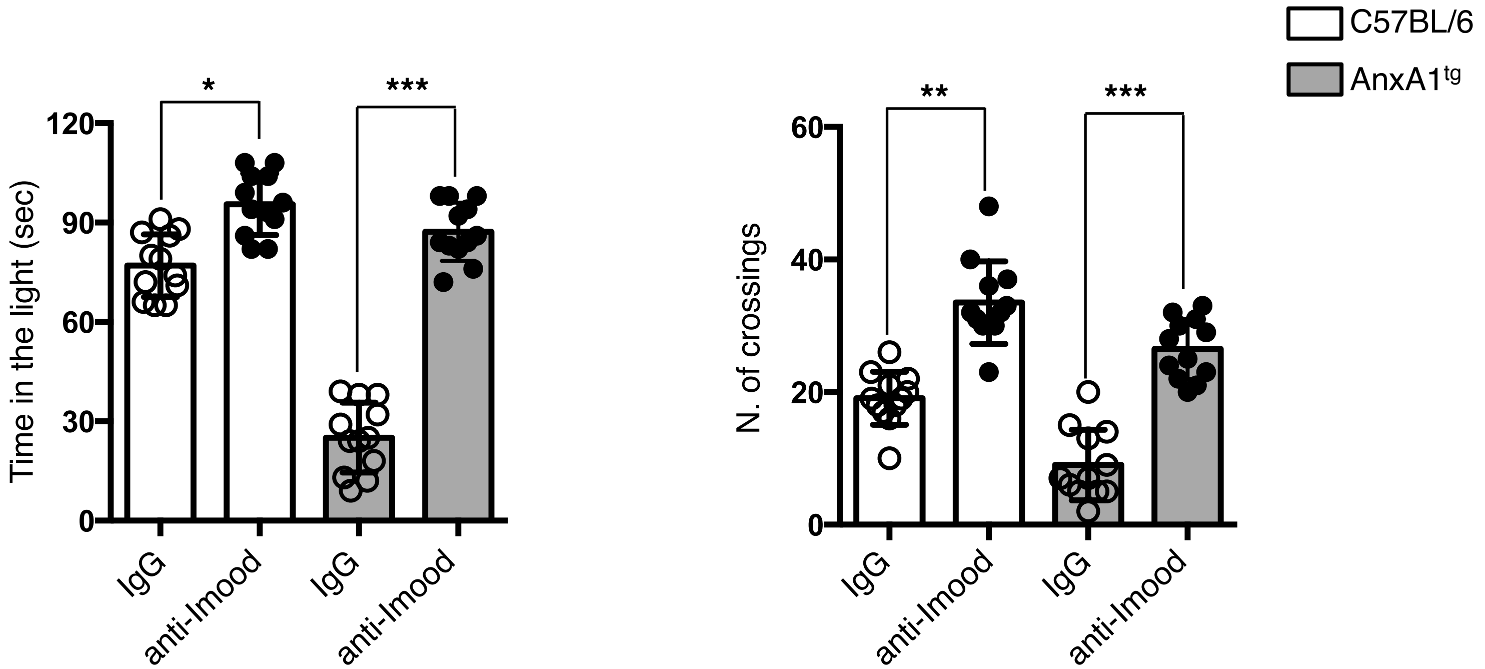

Piras et al., Supplementary Figure 6

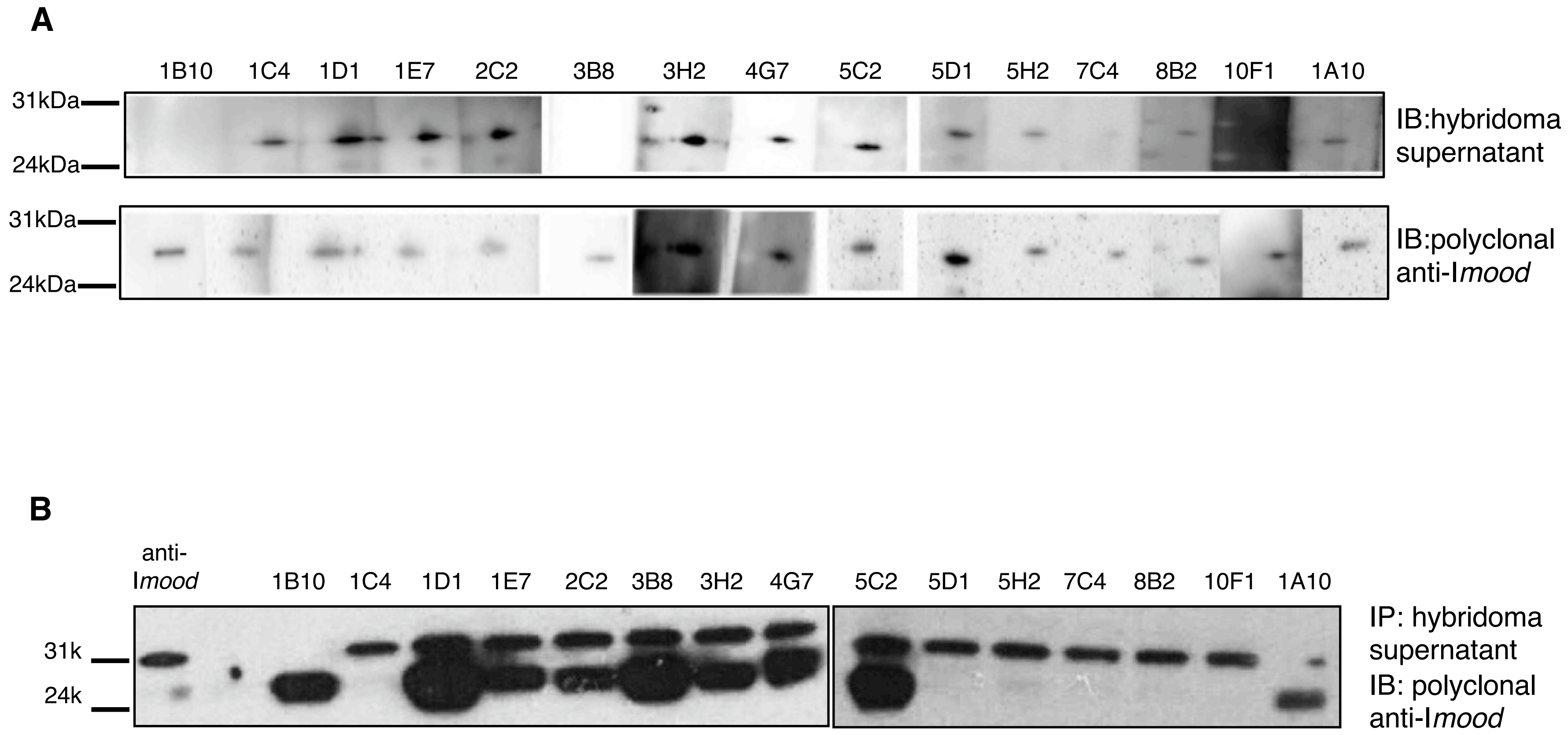
A
1B10 1C4 1D1 1E7 2C2 3B8 3H2 4G7 5C2 5D1 5H2 7C4 8B2 10F1 1A10
31kDa
24kDa
IB:hybridoma supernatant
31kDa
24kDa
IB:polyclonal anti-I*mood*
B
anti-I*mood*
1B10 1C4 1D1 1E7 2C2 3B8 3H2 4G7 5C2 5D1 5H2 7C4 8B2 10F1 1A10
31k
24k
IP: hybridoma supernatant
IB: polyclonal anti-I*mood*

| Gene Symbol | logFC | AveExpr | t | P.Value | adj.P.Val | B | ID |
|---|---|---|---|---|---|---|---|
| Npas2 | 0.557818799 | 5.78134628 | 5.80597972 | 0.00252428 | 0.00252428 | -1.4546163 | 10345675 |
| Il1r2 | -0.390670209 | 6.08537648 | -4.4926299 | 0.00726566 | 0.00726566 | -2.0965173 | 10345752 |
| Il10 | -0.557506558 | 5.70017284 | -4.6127654 | 0.00653896 | 0.00653896 | -2.0269316 | 10349603 |
| Lad1 | -0.375794658 | 7.01663223 | -4.4129303 | 0.0078001 | 0.0078001 | -2.1440528 | 10350159 |
| Rgs16 | -0.860991906 | 9.05322619 | -8.1999755 | 0.0005564 | 0.0005564 | -0.7594912 | 10350733 |
| Ogfrl1 | -0.538326981 | 6.56123977 | -5.0301705 | 0.00459973 | 0.00459973 | -1.8034431 | 10353524 |
| A530032D15Rik | -0.445989488 | 6.26853751 | -4.4840725 | 0.00732094 | 0.00732094 | -2.1015683 | 10356269 |
| Rgs1 | -1.129826065 | 9.46832538 | -9.8333779 | 0.00024446 | 0.00024446 | -0.4849089 | 10358408 |
| Cep350 | 0.428225989 | 8.62697423 | 4.23694216 | 0.0091518 | 0.0091518 | -2.2530168 | 10359078 |
| Slamf7 | -0.788443939 | 7.79162165 | -4.1558183 | 0.00986588 | 0.00986588 | -2.3051493 | 10360173 |
| Atf3 | -1.022463319 | 7.10445479 | -5.6968755 | 0.00273588 | 0.00273588 | -1.4989942 | 10361091 |
| Cd24a | -0.580519819 | 6.56236581 | -4.745834 | 0.00583138 | 0.00583138 | -1.9526719 | 10362896 |
| Gadd45b | -0.497972087 | 9.4067592 | -5.7933151 | 0.00254782 | 0.00254782 | -1.4596952 | 10364950 |
| Chst11 | -0.390488101 | 5.72947422 | -4.5604042 | 0.0068447 | 0.0068447 | -2.0569589 | 10365290 |
| Adgb | 0.465014271 | 5.1336421 | 4.70732705 | 0.00602646 | 0.00602646 | -1.9738623 | 10367805 |
| Zfp365 | -0.389764833 | 5.1036344 | -4.4812411 | 0.00733934 | 0.00733934 | -2.1032423 | 10369783 |
| Appl2 | 0.37277258 | 8.06694601 | 4.47640182 | 0.00737092 | 0.00737092 | -2.1061066 | 10371356 |
| Ublcp1 | -1.55747623 | 5.60590638 | -8.369029 | 0.00050767 | 0.00050767 | -0.7255126 | 10385361 |
| Gm12185 | -0.570191776 | 9.61479565 | -5.6879561 | 0.00275409 | 0.00275409 | -1.5026852 | 10385533 |
| Gm20095 | 1.096913953 | 4.96548787 | 5.58834061 | 0.00296755 | 0.00296755 | -1.5445718 | 10386531 |
| Pygl | -0.44008638 | 5.12593032 | -4.7744666 | 0.00569112 | 0.00569112 | -1.9370706 | 10400844 |
| Enc1 | 0.435954075 | 8.02307288 | 4.32487695 | 0.00844487 | 0.00844487 | -2.1978739 | 10406817 |
| Pxdc1 | -0.363194885 | 6.60267353 | -4.3144408 | 0.00852535 | 0.00852535 | -2.2043449 | 10408629 |
| Hspb1 | 0.527179792 | 6.01710153 | 4.70604363 | 0.0060331 | 0.0060331 | -1.9745727 | 10408928 |
| Nfil3 | -0.507181551 | 6.71353783 | -5.075382 | 0.00443345 | 0.00443345 | -1.7808494 | 10409278 |
| Abhd4 | -0.509341118 | 6.14617759 | -5.2791493 | 0.00376665 | 0.00376665 | -1.6826864 | 10415021 |
| Wdfy2 | 0.547890485 | 6.17079585 | 4.37755989 | 0.00805193 | 0.00805193 | -2.1655064 | 10415818 |
| Snora31 | -0.399120486 | 6.52117287 | -4.4127336 | 0.00780147 | 0.00780147 | -2.1441715 | 10416503 |
| Epsti1 | -0.648756886 | 8.9306251 | -4.4059129 | 0.00784932 | 0.00784932 | -2.1482915 | 10416566 |
| Dzip1 | 0.528984678 | 6.65190054 | 4.75153991 | 0.00580311 | 0.00580311 | -1.9495523 | 10417004 |
| Gm10002 | -0.455973658 | 4.09713874 | -5.3912384 | 0.00345056 | 0.00345056 | -1.6311558 | 10421186 |
| Sec61b | 0.627520977 | 4.75445205 | 5.62633146 | 0.00288391 | 0.00288391 | -1.5284526 | 10426889 |
| Prr13 | -0.443785194 | 8.52102331 | -4.6964368 | 0.00608302 | 0.00608302 | -1.979899 | 10427235 |
| Basp1 | -0.599875859 | 8.17461558 | -4.3124305 | 0.00854095 | 0.00854095 | -2.2055936 | 10427895 |
| Lima1 | -0.406081717 | 6.01865658 | -4.2672713 | 0.00890044 | 0.00890044 | -2.2338385 | 10432540 |
| Ccdc50 | -0.398181446 | 8.53394655 | -4.4172394 | 0.00777005 | 0.00777005 | -2.1414543 | 10434869 |
| Rogdi | -0.421902899 | 8.04589907 | -4.479487 | 0.00735077 | 0.00735077 | -2.10428 | 10437483 |
| Cd86 | -0.564709123 | 6.47923948 | -5.2314581 | 0.00391145 | 0.00391145 | -1.705134 | 10439312 |
| Tigit | -0.608517034 | 9.03770988 | -5.077046 | 0.00442746 | 0.00442746 | -1.7800236 | 10439527 |
| Cdkn1a | -0.949556181 | 7.09920965 | -4.1965583 | 0.00949945 | 0.00949945 | -2.2788159 | 10443463 |
| Hspa1a | -0.51667067 | 5.97955417 | -5.3418057 | 0.00358594 | 0.00358594 | -1.6536714 | 10444589 |
| Armcx3 | -0.731631326 | 3.73855925 | -5.5329516 | 0.00309471 | 0.00309471 | -1.5683979 | 10451851 |
| Ptprs | -0.411202571 | 6.58628295 | -4.3300885 | 0.00840502 | 0.00840502 | -2.1946498 | 10452047 |
| Hmgb1 | -0.811082761 | 5.03683923 | -4.4306742 | 0.00767724 | 0.00767724 | -2.1333736 | 10453252 |
| Klf9 | -0.388433308 | 6.7444597 | -4.4762114 | 0.00737216 | 0.00737216 | -2.1062194 | 10462091 |
| Lgi1 | 0.613641139 | 4.08378219 | 5.9381873 | 0.00229333 | 0.00229333 | -1.4027041 | 10462912 |
| Anxa1 | 2.011582036 | 7.37945593 | 13.464204 | 5.74E-05 | 5.74E-05 | -0.1454681 | 10466606 |
| Il2ra | -0.403079245 | 9.2232664 | -4.7873449 | 0.00562933 | 0.00562933 | -1.9300965 | 10469278 |
| Nostrin | -0.537906308 | 4.48873815 | -5.081942 | 0.00440991 | 0.00440991 | -1.7775962 | 10472514 |
| Ube2l6 | -0.578614856 | 6.10941379 | -6.343464 | 0.00172669 | 0.00172669 | -1.2554432 | 10473356 |
| Olfr1034 | -0.490888519 | 5.45969684 | -5.9729719 | 0.00223679 | 0.00223679 | -1.389375 | 10473494 |
| Olfr1205 | 0.451074555 | 5.92684608 | 4.2357708 | 0.00916167 | 0.00916167 | -2.2537608 | 10473614 |
| Slc12a6 | 0.392343744 | 9.04543389 | 4.50814986 | 0.00716665 | 0.00716665 | -2.0873891 | 10474545 |
| Sord | 0.396654478 | 6.94770547 | 4.5224057 | 0.00707709 | 0.00707709 | -2.0790409 | 10475437 |
| Slc28a2 | -0.47805523 | 7.91275913 | -4.4961192 | 0.00724326 | 0.00724326 | -2.0944614 | 10475487 |
| Dusp2 | -0.662421212 | 8.6986194 | -6.8226934 | 0.00125765 | 0.00125765 | -1.1020306 | 10475782 |
| Cd40 | -0.484384598 | 7.18015415 | -5.6331167 | 0.00286926 | 0.00286926 | -1.5255926 | 10478678 |
| 4930568D16Rik | 0.550698809 | 4.38203959 | 5.6569079 | 0.00281861 | 0.00281861 | -1.5156093 | 10482045 |
| Arhgap11a | -0.732215653 | 6.2250037 | -7.0766047 | 0.00107125 | 0.00107125 | -1.0287831 | 10485963 |
| Atp8b4 | 0.661108766 | 7.38944204 | 5.32893822 | 0.0036222 | 0.0036222 | -1.6595864 | 10487208 |
| F830045P16Rik | -0.377499945 | 4.24595845 | -4.5828211 | 0.0067118 | 0.0067118 | -2.044047 | 10487605 |
| Zfand1 | 0.972463536 | 5.87385827 | 6.97251065 | 0.00114338 | 0.00114338 | -1.0581839 | 10488029 |
| Rpl29 | -0.694456096 | 5.31499455 | -6.0896343 | 0.00205889 | 0.00205889 | -1.3456428 | 10489303 |
| Bmp7 | -0.593626882 | 6.9171321 | -5.2449866 | 0.00386971 | 0.00386971 | -1.6987342 | 10490129 |
| Gapdh | -0.595783578 | 9.63147164 | -4.9303679 | 0.00499354 | 0.00499354 | -1.8544003 | 10492658 |
| Gm12474 | 0.402328529 | 7.45829274 | 4.40615846 | 0.00784759 | 0.00784759 | -2.1481431 | 10494757 |
| Sike1 | 0.464851529 | 7.87249312 | 4.31753868 | 0.00850137 | 0.00850137 | -2.202422 | 10494832 |
| Vmn2r7 | 0.519096086 | 4.15323076 | 5.09048644 | 0.00437948 | 0.00437948 | -1.7733683 | 10498515 |
| Chil3 | -0.487323534 | 4.7609472 | -4.7111853 | 0.00600657 | 0.00600657 | -1.9717282 | 10501020 |
| 2010016I18Rik | 0.463404714 | 8.07437281 | 4.4772655 | 0.00736527 | 0.00736527 | -2.1055951 | 10501048 |
| Agl | 0.390010337 | 5.68076864 | 4.5884604 | 0.00667884 | 0.00667884 | -2.0408122 | 10501699 |
| Gem | -0.770041341 | 6.79431746 | -5.7735494 | 0.00258508 | 0.00258508 | -1.4676596 | 10503334 |
| Hnrnpa3 | 0.403657112 | 8.16274043 | 4.8599001 | 0.0052956 | 0.0052956 | -1.8912962 | 10503370 |
| Penk | -0.809838745 | 7.26116877 | -6.7883958 | 0.0012857 | 0.0012857 | -1.1123308 | 10511363 |
| Gm10588 | 0.429247986 | 5.25831232 | 4.2682392 | 0.00889255 | 0.00889255 | -2.2332292 | 10513004 |
| Speer8-ps1 | 0.581113473 | 6.23396906 | 6.78943185 | 0.00128484 | 0.00128484 | -1.1120182 | 10519811 |
| Lhfpl3 | -0.43348527 | 6.35450975 | -4.5674414 | 0.00680265 | 0.00680265 | -2.0528964 | 10520043 |
| Gtf3c2 | 1.824711665 | 4.87372873 | 8.08093879 | 0.0005941 | 0.0005941 | -0.7843359 | 10529091 |
| Rell1 | -0.447913615 | 7.12784056 | -4.5498298 | 0.00690846 | 0.00690846 | -2.0630793 | 10530130 |
| Tmem176a | -0.794655876 | 7.45954047 | -8.96716 | 0.00037178 | 0.00037178 | -0.616509 | 10538150 |
| Bhlhe40 | -0.57856023 | 7.7033959 | -6.9715721 | 0.00114406 | 0.00114406 | -1.0584529 | 10540472 |
| Cdkn1b | -0.457823151 | 7.00237808 | -4.8648639 | 0.00527363 | 0.00527363 | -1.888672 | 10542317 |
| 8430419L09Rik | 0.509496436 | 8.31398622 | 4.39306535 | 0.00794038 | 0.00794038 | -2.1560744 | 10542340 |
| Tmem176b | -0.861221492 | 8.7625047 | -4.620115 | 0.00649734 | 0.00649734 | -2.0227537 | 10544596 |
| Vmn1r174 | 0.5280242 | 4.52146518 | 6.24163406 | 0.00185168 | 0.00185168 | -1.2908374 | 10550818 |
| Zfp719 | 0.516021502 | 7.22009746 | 6.1322647 | 0.00199811 | 0.00199811 | -1.330028 | 10552440 |
| Lrrc32 | -0.471295227 | 8.12126108 | -4.8655383 | 0.00527066 | 0.00527066 | -1.8883158 | 10555174 |
| Plk1 | -0.542170345 | 7.59122282 | -5.0672225 | 0.00446293 | 0.00446293 | -1.7849047 | 10557156 |
| Mrpl41 | -0.413504215 | 7.0817731 | -4.8292459 | 0.00543368 | 0.00543368 | -1.9075878 | 10561140 |
| Cd22 | -0.38730574 | 5.81290582 | -4.583369 | 0.00670859 | 0.00670859 | -2.0437325 | 10562132 |
| Snord116 | -0.440905479 | 7.57614401 | -4.3155056 | 0.0085171 | 0.0085171 | -2.2036838 | 10564207 |
| 2610034B18Rik | -0.62293187 | 6.41751961 | -7.3588966 | 0.00090135 | 0.00090135 | -0.9531946 | 10564849 |
| Mrpl48 | 0.534638501 | 6.23194814 | 4.95397781 | 0.0048969 | 0.0048969 | -1.8422094 | 10565904 |
| Adam8 | -0.4839193 | 7.44264344 | -5.0991228 | 0.00434897 | 0.00434897 | -1.7691057 | 10568873 |
| Cars | 0.458354339 | 6.89916025 | 4.43296996 | 0.00766151 | 0.00766151 | -2.131996 | 10569458 |
| Dusp4 | -0.677621691 | 7.28271793 | -7.5585278 | 0.00080047 | 0.00080047 | -0.90317 | 10571312 |
| Zfp961 | 0.510381758 | 6.88788166 | 5.93478292 | 0.00229895 | 0.00229895 | -1.4040159 | 10572733 |
| Mt1 | -0.75932243 | 6.42449917 | -5.0598749 | 0.00448967 | 0.00448967 | -1.7885648 | 10574027 |
| Gpr97 | -0.591214167 | 7.05433381 | -4.9771828 | 0.00480406 | 0.00480406 | -1.8303102 | 10574276 |
| Plcg2 | -0.495490201 | 5.83351079 | -4.2543741 | 0.00900633 | 0.00900633 | -2.2419733 | 10575799 |
| Rab4a | 0.660001701 | 6.79503186 | 7.39985008 | 0.00087947 | 0.00087947 | -0.9427087 | 10576391 |
| 2610019F03Rik | 0.52012772 | 7.89015996 | 5.53496162 | 0.00091933 | 0.84208634 | -2.1951276 | 10577226 |
| Usp28 | 0.461535637 | 7.92633842 | 4.75116959 | 0.00580494 | 0.00580494 | -1.9497546 | 10585099 |
| Myo1e | -0.419221101 | 6.1073894 | -5.1025311 | 0.004337 | 0.004337 | -1.7674266 | 10586781 |
| Ccr8 | -0.60881412 | 7.84208574 | -5.9487189 | 0.00227604 | 0.00227604 | -1.3986543 | 10590242 |
| Cxcr5 | -0.687692581 | 5.97790995 | -7.0591529 | 0.00108296 | 0.00108296 | -1.0336529 | 10592888 |
| Ar | 0.395509387 | 6.85958989 | 4.55297288 | 0.00688943 | 0.00688943 | -2.0612581 | 10600901 |
| Erdr1 | -1.165398215 | 10.7662537 | -5.398479 | 0.00343125 | 0.00343125 | -1.6278854 | 10608711 |

| Probe ID | TF4 | TF5 | TF6 | WT4 | WT5 | WT6 | Mean | Mean | Differ | ratio | foldcl | Abs fc | pvalue | Adjusted pvalue | seqna | stran | start | stop | total_ | gene_assignment | mrna_assignment | swiss | unige | GO_b | GO_c | GO_r | pathv | protein_domains |
|---|---|---|---|---|---|---|---|---|---|---|---|---|---|---|---|---|---|---|---|---|---|---|---|---|---|---|---|---|
| 10466606 | 8.36972 | 8.5 | 8.18 | 5.72 | 5.55 | 5.71 | 8.35 | 5.66 | 2.69 | 1.48 | 1.48 | 1.48 | 1.23E-07 | 2.24E-05 | chr19 | - | 20447918 | 20465163 | 25 | NM_010730 // Anxa1 // an | NM_010730 // RefSeq // Mus mu | NM_( | NM_( | NM_( | NM_( | NM_( | NM_( | --- |
| 10418210 | 9.03673 | 9.06 | 8.83 | 9.89 | 9.98 | 9.98 | 8.98 | 9.95 | 0.97 | 0.9 | -1.1 | 1.11 | 1.44E-05 | 0.0026208 | chr14 | - | 27022853 | 27026457 | 39 | NM_025311 // D14Ertd449 | NM_025311 // RefSeq // Mus mu | NM_( | NM_( | NM_( | NM_( | NM_( | --- | --- |
| 10545096 | 6.09332 | 6.19 | 6.99 | 9.07 | 8.67 | 8.89 | 6.42 | 8.87 | 2.45 | 0.72 | -1.4 | 1.38 | 6.08E-05 | 0.0110656 | chr6 | - | 60720390 | 60721053 | 26 | NM_028025 // Mageb16 // | NM_028025 // RefSeq // Mus mu | NM_( | NM_( | NM_( | NM_( | NM_( | --- | --- |
| 10527158 | 9.35675 | 8.95 | 9.27 | 10.1 | 10.3 | 10.2 | 9.19 | 10.2 | 1.01 | 0.9 | -1.1 | 1.11 | 0.000117657 | 0.021413574 | chr5 | + | 143722034 | 143734868 | 25 | NM_007984 // Fscn1 // fas | NM_007984 // RefSeq // Mus mu | NM_( | NM_( | --- | NM_( | NM_( | --- | --- |
| 10393573 | 10.1511 | 10.1 | 10.2 | 9.71 | 9.78 | 9.73 | 10.2 | 9.74 | 0.43 | 1.04 | 1.04 | 1.04 | 0.000165377 | 0.030098614 | chr11 | - | 118254063 | 118263273 | 25 | NM_011150 // Lgals3bp // | NM_011150 // RefSeq // Mus mu | NM_( | NM_( | NM_( | NM_( | NM_( | --- | --- |
| 10574213 | 9.05084 | 8.79 | 8.98 | 9.66 | 9.95 | 9.77 | 8.94 | 9.79 | 0.85 | 0.91 | -1.1 | 1.1 | 0.000167192 | 0.030428944 | chr8 | + | 97269490 | 97275599 | 24 | NM_009137 // Ccl22 // che | NM_009137 // RefSeq // Mus mu | NM_( | NM_( | NM_( | NM_( | NM_( | --- | --- |
| 10360391 | 10.8353 | 10.8 | 10.9 | 10.2 | 10.4 | 10.4 | 10.9 | 10.3 | 0.55 | 1.05 | 1.05 | 1.05 | 0.000220343 | 0.040102426 | chr1 | - | 175850531 | 175872623 | 16 | NM_001045481 // Ifi203 // | NM_001045481 // RefSeq // Mus | NM_( | NM_( | --- | NM_( | NM_( | --- | --- |
| 10577226 | 8.12572 | 8.15 | 8.33 | 7.62 | 7.67 | 7.67 | 8.2 | 7.65 | 0.55 | 1.07 | 1.07 | 1.07 | 0.000225722 | 0.041081404 | chr8 | - | 13952009 | 13974690 | 25 | NM_173744 // 2610019F0 | NM_173744 // RefSeq // Mus mu | NM_1 | NM_1 | NM_1 | NM_1 | NM_1 | --- | --- |

### OCD Patients

#### Sociodemographic Variables

| Variables | All patients |
|---|---|
| Number of patients | 27 |
| Age (years, mean ± SD) | 39.8 ± 14.0 |
| Gender (%) | |
| Male | 44.4 |
| Female | 55.6 |
| Education (%) | |
| Secondary school | 16 |
| High-school | 64 |
| University | 20 |
| Employment (%) | |
| Employed | 40 |
| Unemployed | 48 |
| Student | 4 |
| Retired | 8 |
| Married (%) | 32 |

#### Clinical Variables

| | All patients |
|---|---|
| Age at onset (years, mean ± SD) | 21.8 ± 10.7 |
| Duration of illness (years, mean ± SD) | 18 ± 10.7 |
| Duration of Untreated Illness (months, mean ± SD) | 56.5 ± 72.5 |
| Family history of psychiatric disorder (%) | 72 |
| Psychiatric comorbidity (%) | 64 |
| Drug naive patients | 7.4 |
| Current treatment (%) | |
| Antidepressants | 85.2 |
| Antipsychotics | 55.6 |
| Mood stabilizers | 37 |
| Benzodiazepines | 44.4 |
| Psychotropic compounds (number, mean ± SD) | 2.6 ± 1.5 |
| ≥ 1 psychotropic compounds (%) | 74.1 |
| YBOCS total score (mean ± SD) | 22.6 ± 8.4 |

#### Controls

Age (mean ± SD): 29.2 ± 8.0
Male: 13 (59%)
Female: 9 (41%)

| **Antibodies** | **Clones** | **Source** |
|---|---|---|
| anti-CD3 | clone 145-2C11 | eBioscience |
| anti-CD4 | clone GK 1.5 | eBioscience |
| anti-CD8 | clone 53-6.7 | eBioscience |
| anti-CD25 | clone PC61.5 | eBioscience |
| anti-CD28 | clone 37.51 | eBioscience |
| anti-CD45.1 | clone A20 | eBioscience |
| anti-CD69 | clone H1.2F3 | eBioscience |
| anti-GM-CSF | clone MP1-22E9 | eBioscience |
| anti-IFN-γ | clone XMG1.2 | eBioscience |
| anti-IL10 | clone JES5-16E3 | eBioscience |
| anti-IL-17 | clone eBioTC11-18H10.1 | eBioscience |
| anti-Tdrp | cat. N. NBP1-93675 | Novusbio |
| Alexa Fluor 488- IgG | cat. N. ab150077 | Abcam |
